# An axonal brake on striatal dopamine output by cholinergic interneurons

**DOI:** 10.1101/2024.02.17.580796

**Authors:** Yan-Feng Zhang, Pengwei Luan, Qinbo Qiao, Yiran He, Peter Zatka-Haas, Guofeng Zhang, Michael Z. Lin, Armin Lak, Miao Jing, Edward O. Mann, Stephanie J. Cragg

## Abstract

Depolarisation of distal axons is necessary for neurons to translate somatic action potentials into neurotransmitter release. Studies have shown that striatal cholinergic interneurons (ChIs) can directly drive ectopic action potentials in dopamine (DA) axons and trigger DA release. However, here we show that this action occurs within a broader context of axonal signal integration whereby activation of ChIs and depolarisation of DA axons by nicotinic receptors (nAChRs) limits the subsequent depolarisation and release of DA in response to ensuing activity. We demonstrate that activation of ChIs and nAChRs in *ex vivo* mouse striatum, even when it does not trigger DA release that is detectable by fast-scan cyclic voltammetry, limits DA release for ∼100 ms by depressing subsequent axonal depolarisation and calcium summation. This axonal brake on DA release is stronger in dorsal than ventral striatum, and is unrelated to DA depletion. *In vivo*, antagonism of nAChRs in dorsal striatum elevated extracellular DA levels and promoted conditioned place-preference, underscoring its physiological relevance. Our findings reveal that under physiological conditions *in vivo,* ChIs acting via nAChRs dynamically attenuate DA output driven by DA neuron activity, leading to a predominantly inverse relationship between ACh and DA signalling that varies continuously with ChI activity.

## Introduction

Neuronal axons propagate action potentials and, at vesicular release sites, transform these electrical impulses into neurotransmitter release to activate target cells. Our current understanding of the physiological functions of the neurotransmitter dopamine (DA), such as the encoding of reward prediction error (RPE), has been developed significantly from recordings of action potential firing in the soma of DA neurons^1^. However, mechanisms and inputs acting selectively in or on DA axons are positioned to gate DA release^2-5^, with some *in vivo* evidence that striatal DA release is dissociated from DA somatic activity in some circumstances^6,7^. Tonically active ChIs within striatum constitute only 1-2% of striatal neurons, but arborise densely, and through axo-axonic actions on DA axons can drive short-latency or ‘instantaneous’ DA release events via activation of β2-containing-nAChRs^4,5^ and ectopic axonal generation of action potentials (nAChRs)^8^. However, *ex vivo* experiments have also suggested that when ChIs and DA axons are concurrently stimulated, DA release level might be less than for activation of DA axons alone^2^, suggesting that axonic inputs from ChIs might paradoxically also limit DA release. The ability of ChIs to trigger ectopic action potentials in DA axons might sit within a broader and opposing context of axo-axonic signal integration. Here we test the hypothesis that activation of nAChRs can prevent DA release, by impairing the translation of subsequent action potentials in DA axons into DA release. We find that discrete activation of ChIs and nAChRs which may depolarise DA axons can profoundly and dynamically prevent the ensuing depolarisation of DA axons by incoming activity for up to ∼100 ms. This suppression can occur even for low levels of ChI activation that do not trigger detectable DA release. DA release *in vivo* is then enhanced after antagonism of nAChRs, indicating that a dominant physiological outcome of ChI activity *in vivo* on DA signalling in the intact brain is to operate a dynamically scaling suppression of the amplitude of DA release.

## Results

### Activation of ChIs attenuates subsequent DA release

We first tested the impact of targeted activation of ChIs on subsequent DA release evoked by electrical stimulation. In *ex vivo* striatal slices from ChAT-Cre:Ai32 mice (**Fig. 1a,b**), we used a blue light stimulus (L_stim_) to optogenetically activate ChR2-expressing ChIs to activate nAChRs and drive instantaneous DA release (ChI-driven DA release, DA_ChI_) as previously^4^ followed 8-200 ms later by a second stimulation of a pulse of electrical stimulus (E_stim_) which provides a composite stimulus that drives DA release both directly via activating DA axons (DA_DA_) and indirectly via activation of ChIs (DA_ChI_, which follows DA_DA_ with short latency, ∼10 ms^9,10^). By measuring extracellular DA concentration ([DA]_o_) with fast-scan cyclic voltammetry (FCV), we found in dorsolateral striatum (DLS) (**Fig. 1c,d**) and nucleus accumbens core (NAcc) (**Fig. 1i,j**) that optogenetic stimulation of ChIs evoked instantaneous DA release and then depressed DA release evoked subsequently 8-200 ms later, to ∼15-50% of the [DA]_o_ that could be evoked by an electrical stimulus alone. For intervals ≤ 50 ms in DLS and ≤ 25 ms in NAcc, [DA]_o_ evoked by the second stimulus was significantly less after optogenetic ChI stimulation (when normalised to initial release) than after an initial electrical stimulation in the presence of a β2-nAChR antagonist (DHβE, 1 µM) (**Fig. 1c,d,i,j**). These data suggest that activation of nAChRs can attenuate subsequent DA release over a short interval of up to 100 ms. Note that with nAChRs antagonised, there is a strong inverse relationship between [DA]_o_ and inter-pulse interval for DA axon stimulation, as reported previously^2,11^, whereby this intrinsic short-term depression of DA release is sustained over an extended period of several seconds^2,11^.

**Fig. 1.**
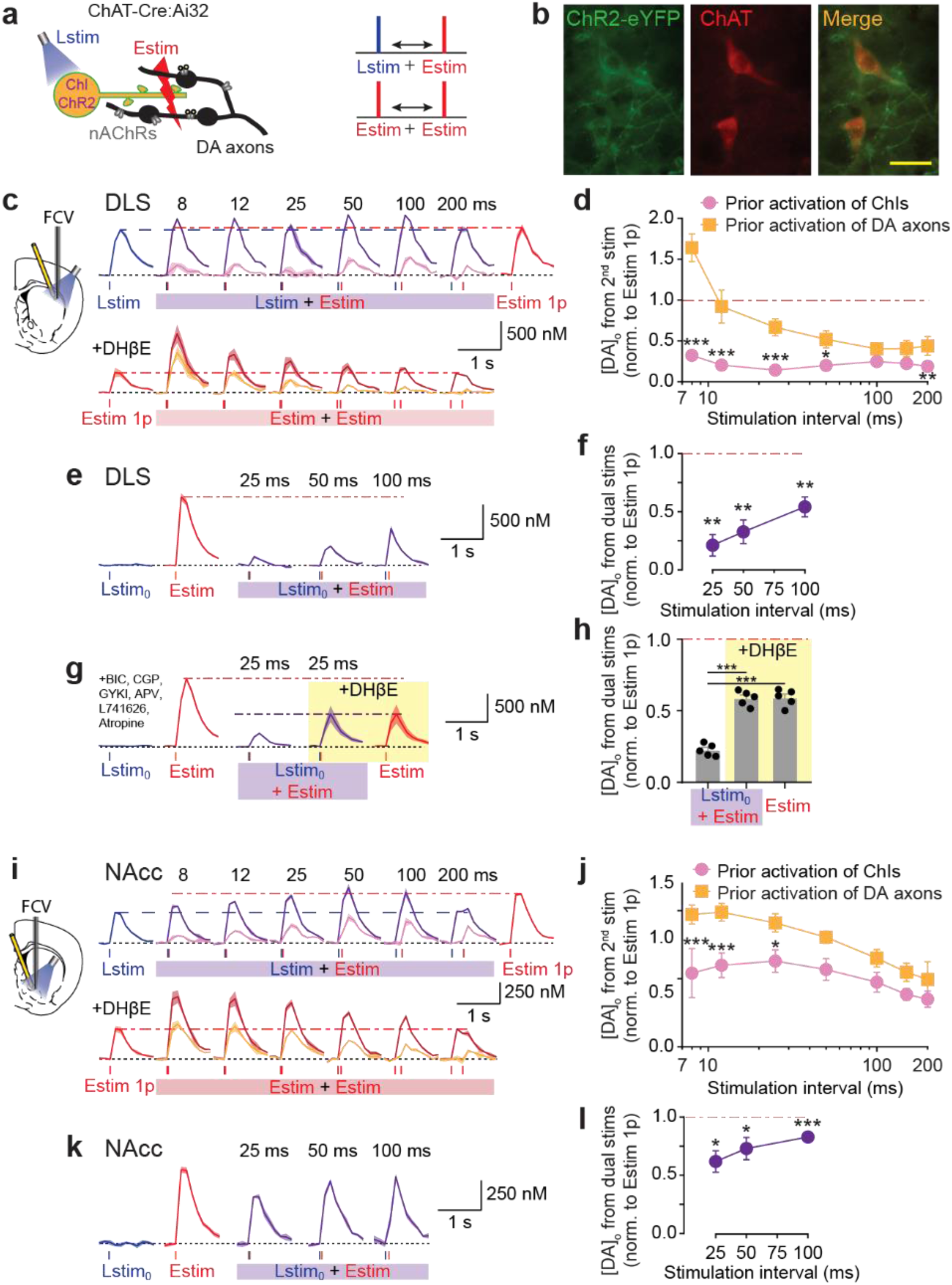
Optogenetic activation of ChIs supresses subsequent DA release evoked by electrical stimulation. **a,** Schematic of stimulation configuration for panels C-J. Blue light stimulation (*Lstim*) of ChR2-eYFP-expressing ChIs, local electrical stimulation (*Estim*) in striatal slices from ChAT-Cre:Ai32 mice. **b,** ChR2-eYP expression in choline acetyltransferase (ChAT)- immunoreactive striatal neurons. Scale bar, 40 µm. **c,i,** *Upper*, mean transients from representative experiments of [DA]_o_ (± SEM) evoked by a single pulse of Lstim (*blue lines*), or Estim (*red lines*), or the composite response to paired Lstim plus Estim pulses (*purple*) at ISIs of 8-200 ms in DLS (**c**) and NAcc (**i**). *Pink trace*, [DA]_o_ attributable to the paired Estim after subtraction of [DA]_o_ due to Lstim. *Lower,* mean transients of [DA]_o_ (± SEM) evoked by single or paired electrical pulses (*red lines*) in the presence of DHβE (1 µM) in DLS **(c)** and NAcc **(i)**. *Orange* trace, [DA]_o_ attributable to the paired Estim after subtraction of [DA]_o_ due to single Estim. **d,j,** Mean peak [DA]_o_ (± SEM) evoked by the paired Estim normalised to [DA]_o_ evoked by a single Estim, versus ISI in DLS (**d**, N = 5 animals) and NAcc (**j**, N = 5 animals). *P < 0.05, **P < 0.01, ***P < 0.001, Two-way ANOVA with Fisher’s LSD test *post hoc*. **(e,g,k)** Mean transients from representative experiments of [DA]_o_ (± SEM) evoked by a subthreshold light pulse (*Lstim_0_*, *blue*), single Estim (*red*), or paired Lstim and Estim (*purple*) at ISIs of 25 - 100 ms in DLS (**e,g**) or NAcc (**k**). **f,h,l,** Mean peak [DA]_o_ (± SEM) evoked by the dual stimuli normalised to [DA]_o_ evoked by a single Estim, versus ISI in DLS (**f,h**, N = 5 animals) and NAcc (**l**, N = 5 animals), with antagonists of GABA_A_, GABA_B_, AMPA, NMDA, D_2_, and mAChRs presence and compared to after nAChRs were blocked (**g**,**h**). ***P < 0.001, t-test versus single electrical stimulation.

We explored whether the greater depression of subsequent DA release seen after optogenetic ChI activation than after electrical stimulation of DA axons in the presence of an nAChR antagonist was due to depletion of the DA vesicle pool^9^. A low intensity of light (Lstim_0_) (**Extended Data Fig. 1**) was applied to stimulate ChIs at a minimal level for which no instantaneous [DA]_o_ could be detected by FCV (equivalent to < 0.5% of DA_ChI_ evoked by normal Lstim) (**Fig. 1e,k**). This Lstim_0_ stimulus will ensure that > 99.5% of the DA release pool is still available for release. However, we found that even using Lstim_0_ to stimulate ChIs, the [DA]_o_ evoked by a subsequent electrical pulse was depressed to as little as 20% in DLS and 60% in NAcc of the [DA]_o_ evoked by an electrical pulse alone (**Fig. 1e,f,k,l**). The depression of DA release after prior ChI activation cannot then solely be explained by DA depletion, indicating that activation of ChIs/nAChRs limits subsequent DA release through a mechanism independent from DA vesicle pool availability.

We tested whether the depression of DA release was due to the initial Lstim_0_ of ChIs causing depletion of ACh, and so compromising ACh available to drive DA_ChI_ at the subsequent Estim (which is made up of DA_DA_ and DA_ChI_). We prevented DA_ChI_ altogether, by using the nAChR antagonist DHβE, and repeated the experiment to test two competing hypotheses. If the low [DA]_o_ evoked by Estim (DA_DA_ + DA_ChI_) after Lstim_0_ was due to a loss of DA_ChI_, then the [DA]_o_ evoked by Estim when nAChRs are antagonised (DA_DA_) might reach a similar but never a greater value. Conversely, if Lstim_0_ of ChIs reduces subsequent DA release evoked by Estim (DA_DA_ + DA_ChI_), then [DA]_o_ evoked by Estim when nAChRs are antagonised (DA_DA_) might be able to exceed this level of release. Testing these hypotheses in DLS, we found that the DA_DA_ evoked by Estim after Lstim_0_ was indeed significantly higher when nAChRs were antagonised than not (**Fig. 1g,h**) (and was equivalent to [DA]_o_ evoked by Estim when nAChRs were antagonised in the absence of prior Lstim_0_), indicating that subthreshold stimulation of ChIs actively attenuates subsequent DA release, including DA_DA_. These experiments were conducted in the presence of a cocktail of antagonists for GABA_A_ and GABA_B_ receptors (bicuculline 10 μM, 4 μM CGP 55845), AMPA and NMDA glutamate receptors (10 µM GYKI 5246, 50 µM D-APV), D2 receptors (1 µM L-741,626), and mAChRs (2 µM atropine) (**Fig. 1 g,h**), therefore allowing us to exclude the possibility that ChI-dependent attenuation of DA release was mediated by indirect activation of these other candidate receptors and transmitter networks.

To demonstrate further that stimulation of ChIs depresses subsequent DA_DA_ release, (and to further avoid confounding effects on presynaptic short-term plasticity of DA or ACh release probabilities resulting from summation of stimuli), we used a dual optogenetic approach in double transgenic ChAT-Cre:DAT-Cre mice to tailor stimulation to ChIs versus DA axons, in DLS and NAcc. ChR2 packaged in AAV2 (for anterograde axonal transport)^12^ was injected to the striatum for preferential expression in ChIs and to minimise expression in DA axons (**Fig. 2a,b**). Chrimson (packaged in AAV5) was injected in SNc/VTA for expression in DA neurons and axons (**Fig. 2a,b**). In striatum, we used a minimal intensity of blue light (480 nm laser, 2 ms) to preferentially stimulate ChIs without driving detectable [DA]_o_, and a threshold level orange light (585 ± 22 nm LED, 2 ms) to preferentially stimulate DA axons and drive detectable [DA]_o_ (DA_DA_) (**Fig. 2c,e, Extended Data Fig. 2**), in the presence of a cocktail of antagonists for GABA, AMPA, NMDA, D_2_ and mAChR receptors. After minimal blue light stimulation of ChIs, [DA]_o_ evoked by subsequent orange light stimulation of DA axons 25 ms afterwards was significantly lower compared to [DA]_o_ evoked by an orange light pulse alone, in both DLS (**Fig. 2c,d**) and NAcc (**Fig 2e,f**). Furthermore, this attenuation of light-evoked [DA]_o_ by prior ChI stimulation was prevented by antagonising nAChRs with DHβE (**Fig. 2c,d,e,f**). Therefore, activation of nAChRs at a level below the threshold for driving DA release can profoundly depress subsequent DA_DA_ through a mechanism unrelated to changes to availability of DA vesicles.

**Fig. 2.**
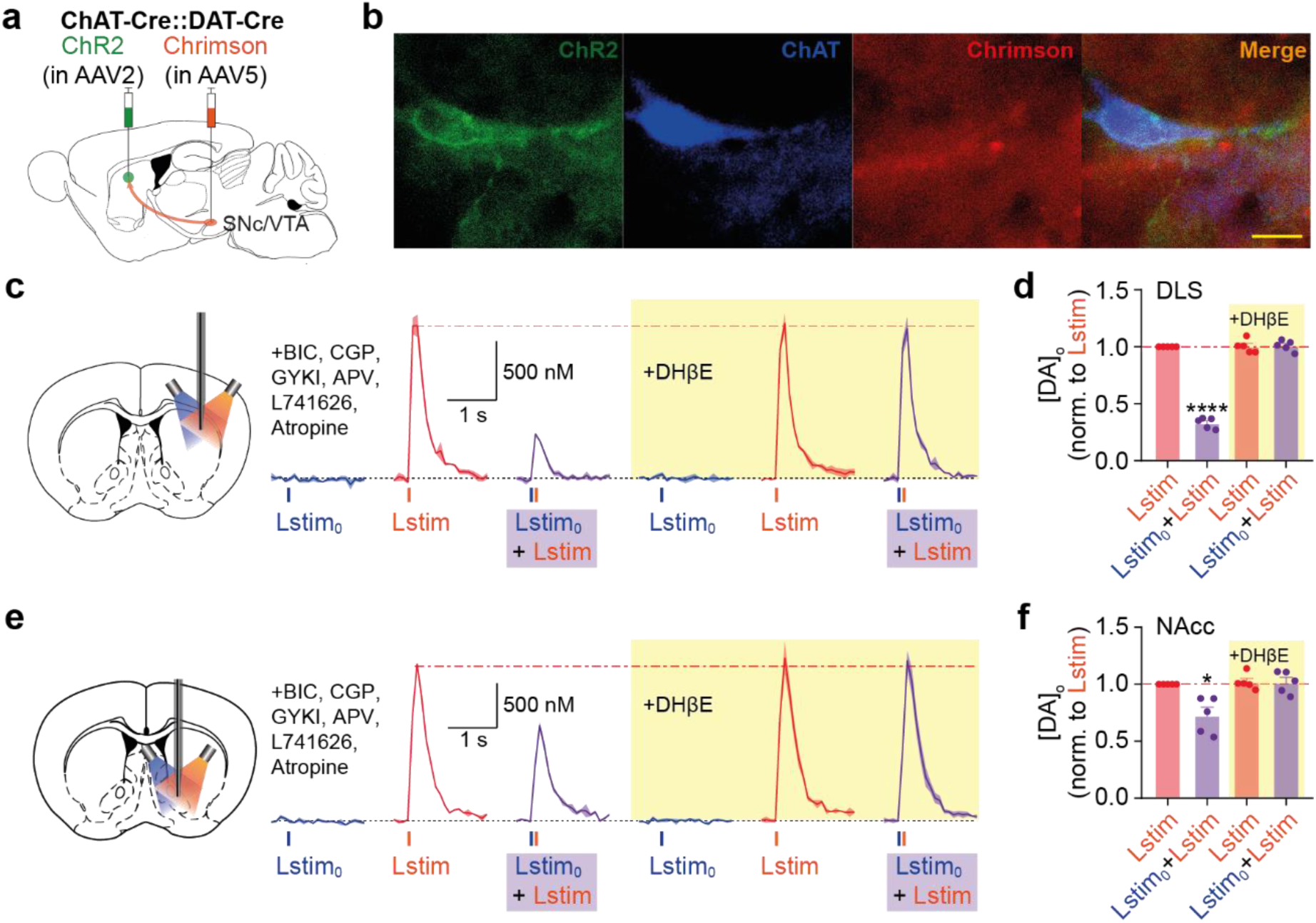
A dual optogenetic approach shows that subthreshold activation of ChIs supresses subsequent DA release. **a,** Schematic of virus injection in double transgenic ChAT-Cre:DAT-Cre mice. **b**, Images of a ChI with expression of ChR2 (green) co-labelled with ChAT-immunoreactivity (blue), alongside Chrimson positive (red) DA axons (scale bar, 20 µm). **c**,**e**, Subthreshold blue light (*Lstim_0_*) stimulation to preferentially activate ChR2-eYFP-expressing ChIs, orange light stimulation to preferentially activate Chrimson-expressing DA axons to drive DA release, in striatal slices in DLS (**c**) or NAcc (**e**). Mean transients of [DA]_o_ (± SEM) from representative experiments evoked by a single pulse of blue Lstim_0_ (*blue lines*), or orange Lstim (orange *lines*), or the response to paired blue Lstim_0_ plus orange Lstim (*in purple shade*) at an interval of 25 ms, in the presence of receptors antagonists for GABA_A_, GABA_B_, AMPA, NMDA, D_2_, and mAChRs. Yellow shading indicates the presence of DHβE (1 µM). **d,f,** Mean peak [DA]_o_ (± SEM) evoked by each stimulus paradigm normalised to [DA]_o_ evoked by a single orange Lstim in DLS (**d**, n = 5 in N = 3 animals) and NAcc (**f**, n = 5 in N = 4 animals). *P < 0.05, ****P < 0.0001, One-sample t-test versus single orange Lstim before adding DHβE.

We then used an alternative stimulation paradigm to characterise the time course of ChI-dependent depression of subsequent DA_DA_ release. We combined an initial electrical stimulus, which drives DA_DA_ + DA_ChI_, with a subsequent targeted optogenetic stimulation of DA_DA_, by activating DA axons expressing ChR2 with blue light in *ex vivo* striatal slices from DAT-Cre mice (**Fig. 3a,b**) in DLS (**Fig. 3c,d**) and NAcc (**Fig. 3g,h**) and examined the difference in depression seen when β2-nAChRs were available versus antagonised (DHβE, 1 µM). [DA]_o_ evoked by the subsequent light pulse was depressed compared to [DA]_o_ evoked by a single light pulse alone and for inter-stimulus intervals up to ∼100 ms (DLS) or ∼50 ms (NAcc). This depression was relieved when nAChRs were antagonised. The relief from depression was not a consequence of less DA depletion arising from the lower levels of DA release evoked by initial electrical stimulation: in wild-type mice, we halved the level of initial DA release to a level similar to that seen for a single pulse in the presence of DHβE, by lowering electrical stimulation intensity (Estim_50_), but did not find a corresponding increase in the [DA]_o_ evoked by a subsequent full intensity electrical stimulation, which evoked the same [DA]_o_ as following a full strength stimulus in DLS (**Fig. 3 e,f**), and only slightly greater [DA]_o_ in NAcc (**Fig. 3 i,j**). Rather, activation of nAChRs by ChIs depresses DA release during subsequent activation of DA axons through a mechanism independent of prior ACh or DA release, that is more pronounced and longer lasting in DLS than NAcc, for durations that are notably similar to the duration of ChI pauses.

**Fig. 3.**
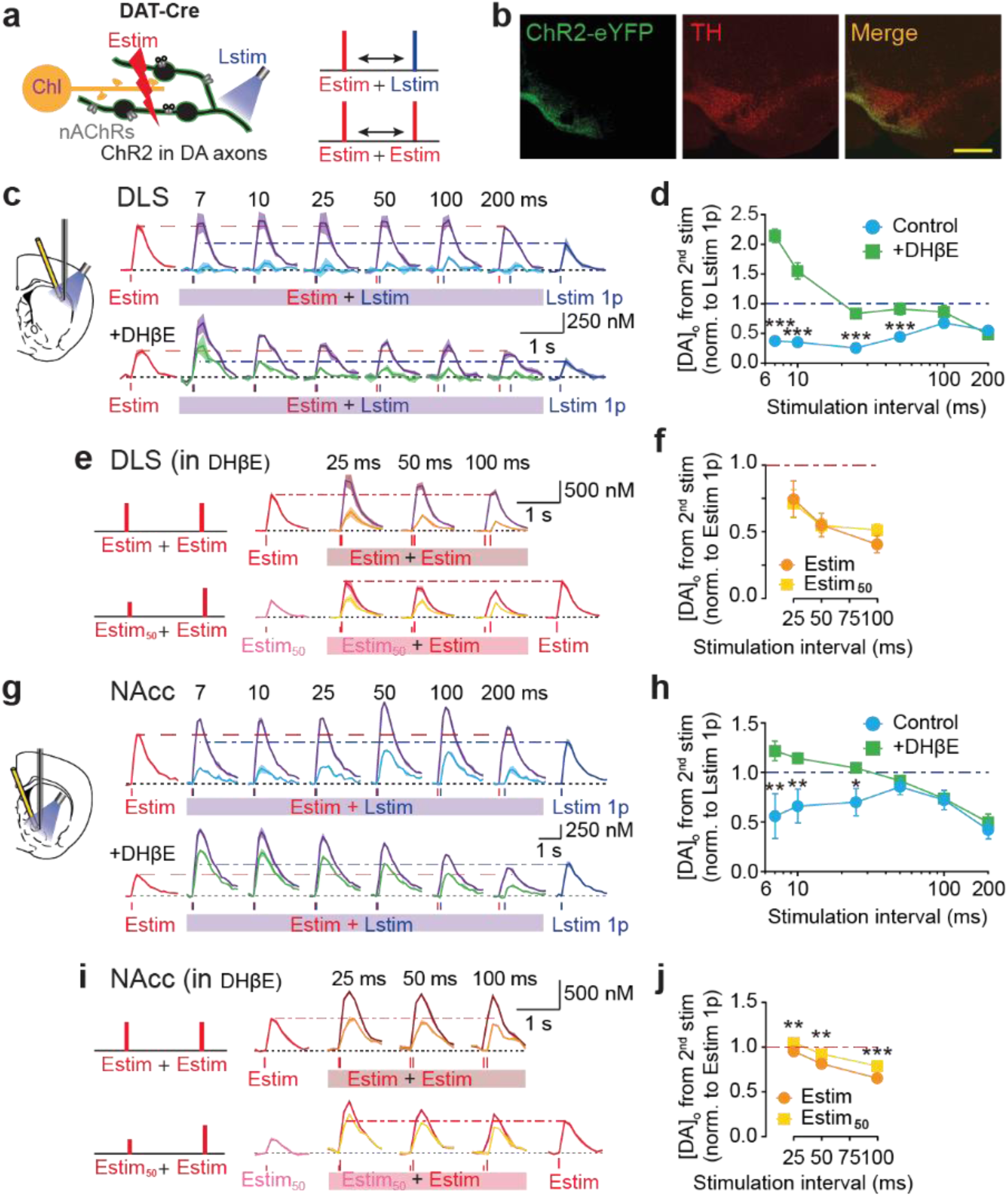
Activation of ChIs depresses subsequent DA release evoked by optogenetic stimulation. **a,** Schematic of stimulation configuration for panels **c**-**j**. Local electrical stimulation (*Estim*) in striatal slices and blue light stimulation (*Lstim*) of ChR2-eYFP-expressing DA axons. **b,** ChR2-eYFP expression in midbrain DA neurons co-immunoreactive for DAT in DAT-Cre mice, after example VTA injection. Scale bar, 400 µm. **c,g,** Mean transients from representative experiments of [DA]_o_ (± SEM) evoked by a single pulse of Estim (*red lines*) or Lstim (*blue lines*), or paired Estim plus Lstim pulses (*purple*) at ISI of 7 - 200 ms in DLS (**c**) and NAcc (**g**) when (*upper*) nAChRs can be active (no DHβE), or (lower) when nAChRs are antagonised (DHβE present) in DAT-Cre mice. *Light blue* trace, [DA]_o_ attributable to the paired Lstim after subtraction of [DA]_o_ due to Estim without DHβE, *green* trace, [DA]_o_ attributable to the paired Lstim in the presence of DHβE. **d,h,** Mean peak [DA]_o_ (± SEM) evoked by the paired Lstim normalised to [DA]_o_ evoked by a single Lstim, versus ISI in DLS (**d**, N = 5 animals) and NAcc (**h**, N = 5 animals). *P < 0.05, **P < 0.01, ***P < 0.001, Two-way ANOVA with Fisher’s LSD test *post hoc*. **e,i,** Mean transients from representative experiments of [DA]_o_ (± SEM) evoked by a single full strength Estim (*red*), paired Estims (*brown*), or a single low intensity Estim (Estim_50_, *pink*) with paired full-strength Estim (*dark red*) at ISIs of 25 - 100 ms in DLS (e) and NAcc (**I**) in wild-type animals. *Orange* and *yellow* traces, [DA]_o_ attributable to the paired Estims after subtraction of [DA]_o_ due to Estim 1p. **f,j,** Mean peak [DA]_o_ (± SEM) evoked by the paired stimulations versus ISI in DLS (f, N = 5 animals) and NAcc (j, N = 5 animals). DHβE was present in **e**, **f**, **i**, and **j**. **P < 0.01, ***P < 0.001, Two-way ANOVA with Fisher’s LSD test *post hoc*.

### Activation of nAChRs limits subsequent increase in axonal Ca^2^ and membrane depolarisation

DA release is strongly governed by axonal activation mechanisms upstream of Ca^2+^ entry, as well as those governing Ca^2+^ entry, intracellular buffering and signalling^13-16^. To understand the depression of DA release that follows initial activation of striatal nAChRs, we first tested whether nAChR activation also limited subsequent axonal Ca^2+^ influx at successive stimuli. In brain slices of DAT-Cre:Ai95D mice (**Fig. 4a,b**), we imaged Ca^2+^ reporter GCaMP6f in DA axons in DLS as previously^14^, following single versus 4-pulse electrical stimuli (100 Hz, or 143 Hz corresponding to 7ms IPI), with and without nAChR antagonism. DA release evoked by this protocol is very sensitive to nAChR activation^2,17^: the ratio of [DA]_o_ evoked by a 100 Hz pulse train versus single pulses is only a little over 1 when nAChRs are activated but is markedly increased when nAChRs are antagonised. In parallel, the ratio of axonal GCaMP6f fluorescence for a train compared to a single pulse was only slightly greater than 1 when nAChRs were active (**Fig. 4c,d, Extended Data Fig. 4**), but markedly greater than 1 when we antagonised β2-nAChRs (DHβE, 1 µM) (**Fig. 4c,d, Extended Data Fig. 4**), indicating that nAChR activation limits summation of axonal [Ca^2+^]_i_ during subsequent stimuli. Absolute levels were not compared before and after DHβE because of the propensity for signal decay/bleaching over time and the non-linear relationships between [Ca^2+^]_o_ and neurotransmitter release^18^.

**Fig. 4.**
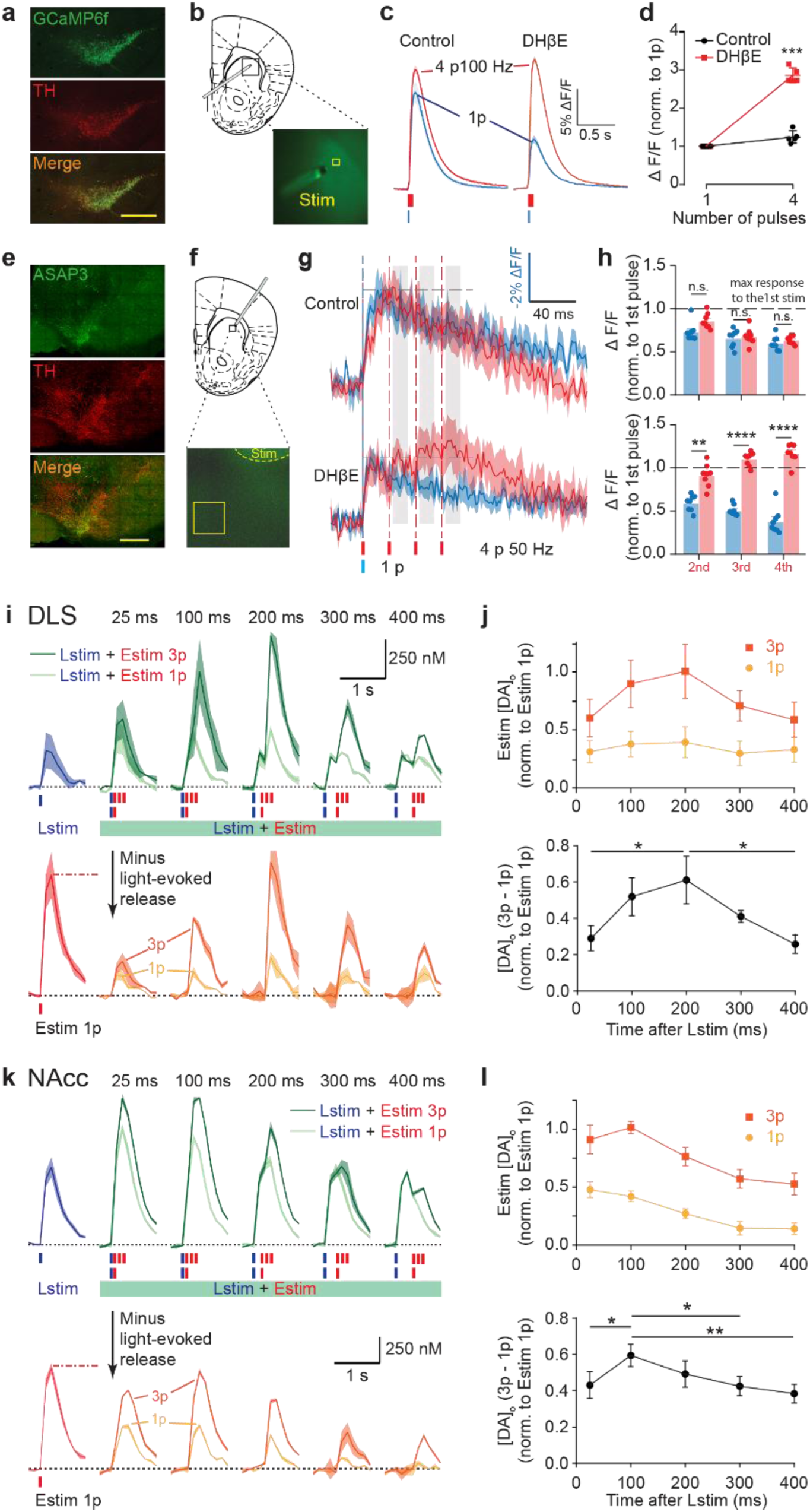
Activation of ChIs and nAChRs attenuates repetitive axonal depolarisation and calcium summation in DA axons, and at longer intervals limits nAChR control of DA release. **a**, Images of VTA and SNc from DAT-Cre:Ai95D mice showing GCaMP6f-eGFP expression (green) in TH-immunopositive neurons (red). **b,** Illustration of stimulation configurations in live tissue. **c,** Examples of Ca^2+^-imaging responses (changes to GCaMP6f fluorescence, ΔF/F) (mean ± SEM from duplicates) in a DA axon population imaged in DLS in response to single or trains of 4 electrical pulses (4p) at 100 Hz in control conditions (*left*) and in the presence of DHβE (1 μM) (*right*). **d,** Mean peak values (± SEM) for GCaMP6f ΔF/F vs. pulse numbers. Data are normalised to value for 1 pulse (N = 5 animals). *** P < 0.0001 paired t-test [DA]_o_ from 4 pulses of stimulation. **e**, Images of VTA and SNc from DAT-Cre mice with ASAP3 expression (green) and TH-positive neurons (red). **f**, Illustration of stimulation configuration in live tissue. **g**, Averaged transients of voltage sensor (changes to ASAP3 fluorescence, - ΔF/F, mean ± SEM) in striatal DA axons to single (blue) or 4 electrical pulses at 50 Hz (red) before and after antagonising nAChRs with DHβE. Scale bar vertical axis applies to 1p data only. For the 4-pulse data, the peak value seen after the first pulse of stimulation is scaled to match the peak of transients seen for single pulses (n = 7 recordings in N = 4 animals). **h**, The averaged responses at the time after the successive pulses in the pulse train (grey in **g**) before and after DHβE. **i,k,** Mean transients from representative experiments of [DA]_o_ (± SEM) evoked by a single pulse of Lstim (*blue*) or Estim (*red lines*), or Lstim paired with either 1 or 3 Estim pulses (*green*) at ISIs of 25 - 400 ms spanning activation, desensitisation, and resensitisation in DLS (**i**) and NAcc (**k**) in striatum of ChAT-Cre: Ai32 mice. *Yellow and orange* traces, [DA]_o_ attributable to the paired stimuli after subtraction of Lstim. j,l, *Upper*, mean peak [DA]_o_ (± SEM) evoked by paired Estims for 1p (*yellow*) and 3p (*orange*) normalised to [DA]_o_ evoked by a single Estim, versus ISI, for DLS (**j**, N = 5 animals) and NAcc (**l**, N = 5 animals). *Lower*, differences between [DA]_o_ evoked by 3p and 1p Estims. *P < 0.05, **P < 0.01, One-way ANOVA with Tukey test *post hoc*.

To test whether nAChRs directly limit axonal depolarisation during successive stimulation of DA axons, we imaged the ASAP3 voltage sensor^19^ expressed in DA axons in DLS in brain slices following single versus 4-pulse electrical stimuli (50 Hz), with and without nAChR antagonism (**Fig. 4e,f, Extended Data Fig. 3**). In the absence of a nAChR antagonist, depolarization of DA axons could be detected by the first but not subsequent stimulus pulses in the train (**Fig. 4g,d**), whereas when nAChRs were antagonised, depolarization of DA axons was detected for each and every pulse (**Fig. 4g,h**). These results indicate that activation of nAChRs initially depolarizes DA axons, in agreement with other reports^20^, but in consequence, for a time window of ∼50-100 ms this leads to an extended refractory-like period, when subsequent depolarisation and Ca^2+^ entry are limited, with further DA release prevented.

### nAChRs activated by initial excitation are off during ChI rebound

In an *in vivo* multiphasic response, ChIs show burst-pause activity followed by a ‘rebound’ increase in activity some ∼100-300 ms after initial excitation^21,22^. At these intervals, the ChI-dependent depression of DA release from the initial excitation will have dissipated (see Fig. 1). However, for the ensuing ∼1 second, it has been shown that further DA release is resistant to targeted optogenetic stimulation of ChIs but not DA axons^4,9^. This selective refractoriness of DA release to activation of ChIs but not DA axons suggests that nAChRs cannot be activated by ACh at the time of rebound ChI activity. One potential explanation could be the desensitisation of nAChRs that follows their activation^23-25^ and has been suspected to play a role in DA release dynamics during stimulus trains^26^. We tested whether the DA release dynamics at the timepoints of ChI rebound activity are consistent with lack of nAChR activation e.g. by nAChR desensitisation, by exploiting the observations that whereas the activation of nAChRs limits DA release at subsequent pulses within 100 ms (as in Fig. 1), the desensitisation or inactivation of nAChRs allows subsequent release during a high frequency train and increases the ratio of [DA]_o_ evoked by trains versus single pulses^2^. In striatal slices from ChAT-Cre:Ai32 mice, we light-activated ChIs, then 25-400 ms later, applied either single or triplets of electrical pulses (100 Hz) to explore how the difference in [DA]_o_ evoked by triplet versus single pulses (100 Hz) varied with time (**Fig. 4i,k**, also see **Extended Data Fig. 3C**). The difference in [DA]_o_ evoked by triplet versus single pulses varied with interval, peaking at ∼200 ms in DLS and ∼100 ms in NAcc, and decaying by 400 ms (**Fig. 4i-l**). Neither muscarinic nor D_2_-receptor activation were responsible for these dynamics (**Extended Data Fig. 3d,e).** These dynamics likely reflect the initial inability and then renewed ability to activate nAChRs, which could reflect the timecourse of nAChR desensitisation and resensitisation. They indicate that nAChRs will not be in an activatable state at ChI rebound activity ∼100-300 ms after initial excitation, and will not then be strongly limiting or driving DA release at this interval. The regional differences in timecourses parallels differences in α4β2-nAChR stoichiometries^27,28^ which might contribute to different desensitisation time courses.

### nAChR antagonism in vivo promotes DA release, DA axon activity and induces conditioned place-preference

*In vivo,* ChIs and DA neurons fire tonically at 3 – 10 Hz, which can lead to a balance of ongoing nAChR activation and desensitisation that will govern net impact on DA. Desensitisation of nAChRs might continuously limit how nAChRs suppress DA output. We tested *in vivo* whether nAChRs are able to depress DA release or are desensitised. In wild-type urethane-anesthetised mice, we found that [DA]_o_ evoked in DLS by local electrical stimulation with brief pulse trains and detected using FCV was increased following systemic injection of nAChR antagonist mecamylamine (**Fig. 5a-d**) indicating that after ChI activation, nAChR activation is limiting DA release *in vivo*. Previous work in NAcc has shown that systemic nAChR antagonist also increases DA release in response to reward, in freely moving rats^29^.

**Fig. 5.**
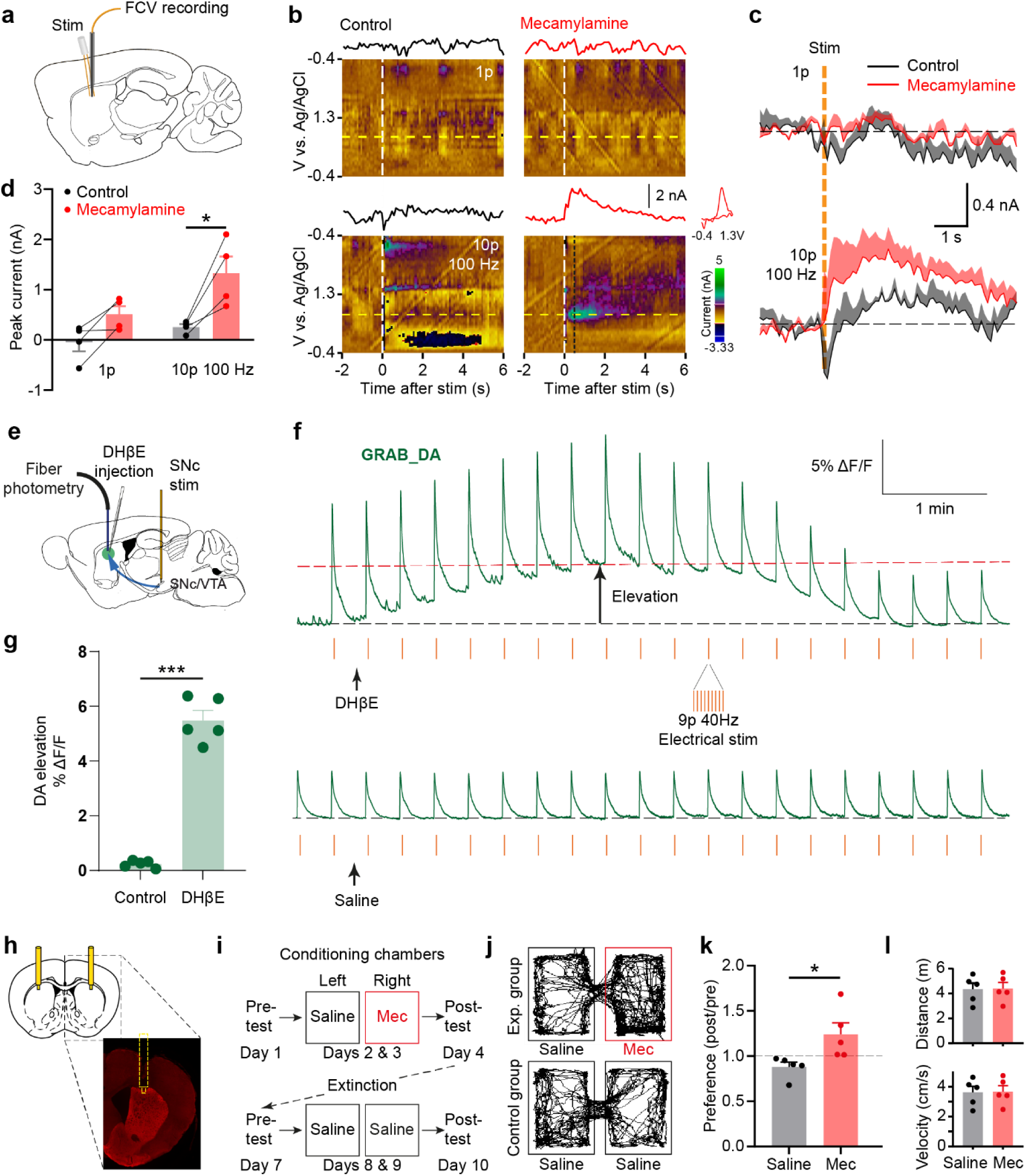
nAChR antagonism in DLS *in vivo* promotes DA release and conditioned place preference. **a,** Schematic of configuration of FCV recording and stimulation electrodes in DLS. **b,** Colour plots and representative line plots of oxidation current (from yellow dashed row at approximately +0.7 V) of voltammetric DA current vs. time and a corresponding DA cyclic voltammogram (from timepoint of black dotted line) before and after mecamylamine (i.p.). **c,** DA signals (mean+SEM) evoked by 1 pulse or 10 pulses at 100 Hz before (*black*) and after (*red*) mecamylamine (2 mg/kg i.p.). **d,** Mean peak DA currents (± SEM). **P < 0.01 for Sidak’s post-hoc multiple comparisons test (N = 4 animals). **e**, Schematic of configuration of fibre photometry recording in DLS and stimulation in SNc. **f**, Example traces of fibre photometry recording of GRAB-DA_2m_ signal with local injection of DHβE (0.07 µg in 200 nl saline) or saline (200 nl) showing an elevation in [DA]_o_ tone after DHβE. Intermittent electrical stimulation (9 pulses at 40 Hz, red lines) of SNc was applied at 0.1 Hz as a reference. **g**, Mean peak change in tonic DA (± SEM) after local injection of DHβE or saline control. ***P < 0.001 paired t-test (n = 5 from N = 3 animals). **h,** Schematic illustration of a bilateral cannula system for local infusion to dorsal striatum, and an example hemi-slice with DAPI staining. **i**, Conditioning paradigm. Mice were conditioned for 4 times over two days (20 min per session) after local infusion of mecamylamine (10 µg/side, right chamber, *red*) or saline (0.5 µl over 1 min, left chamber, *black*), whereas the control groups received saline for both chambers. **j,** Representative tracking traces from the post-conditioning day. **k,** Preference for the right chamber after conditioning. * P < 0.05 paired t-test, N = 5 animals in each test. **l**, Total travel distances and velocity of movements in open-field test (N = 5 animals).

Furthermore, using fibre photometry in urethane-anaesthetised mice, we identified the impact of local antagonism of β2-nAChRs in DLS on the tonic level of DA release detected using fluorescent sensor GRAB-DA2m^30^in wild-type mice, and separately, on DA axon activity reported by GCaMP6f in DAT-Cre:Ai95 mice. We gained a reference amplitude for these changes to DA tone by comparing to a reference signal evoked by electrical stimulation of midbrain (9p, 40 Hz every 10 s). Note that evoked signal amplitudes were not analysed owing to bleaching over time that led to uncorrected signal rundown. Local striatal injection of DHβE (0.07 µg in 200 nl) significantly increased the non-evoked fluorescence of GRAB-DA_2m_ (**Fig. 5e-g**) and GCaMP6f (**Extended Data Fig. 5**) compared to saline controls, by amplitudes comparable to those evoked by midbrain stimulation. Therefore, *in vivo,* ChIs acting at nAChRs operate a suppression of DA axon activity and DA release.

We then tested in freely moving mice whether nAChRs in dorsal striatum could modify reward-related learning as might be predicted for modulation of DA function^31^. After two days of daily conditioning, mice developed a conditioned-place preference for the chamber conditioned with intrastriatal diffusion in DLS of nAChR antagonist mecamylamine but not saline controls (**Fig. 5h-l**). Our findings are consistent with *in vivo* studies in adjacent NAc showing that nAChR antagonists increase reward-evoked DA levels and promote reward-related learning^29,32^.

### Tonic and multiphasic activity in ChIs depresses DA release in a computational model

We developed a computational model to predict how nAChRs impact on [DA]_o_ *in vivo* in DLS and NAcc during the dynamic and multiphasic activity in ChIs that is coincident with phasic activity in DA neurons (**Fig. 6a**). The model incorporates the timings of the dynamic suppression of DA release by ChIs (from Fig. 3), and the decay (desensitisation) and recovery (resensitisation) of nAChR control of DA release (from Fig. 4), as well as the extracellular kinetics of DA signals (detected after electrical stimulation when nAChRs are off, Fig. 1). We validated that the model could simulate our *ex vivo* observations of Fig. 1 d,j (**Extended Data Fig. 4**).

**Fig. 6.**
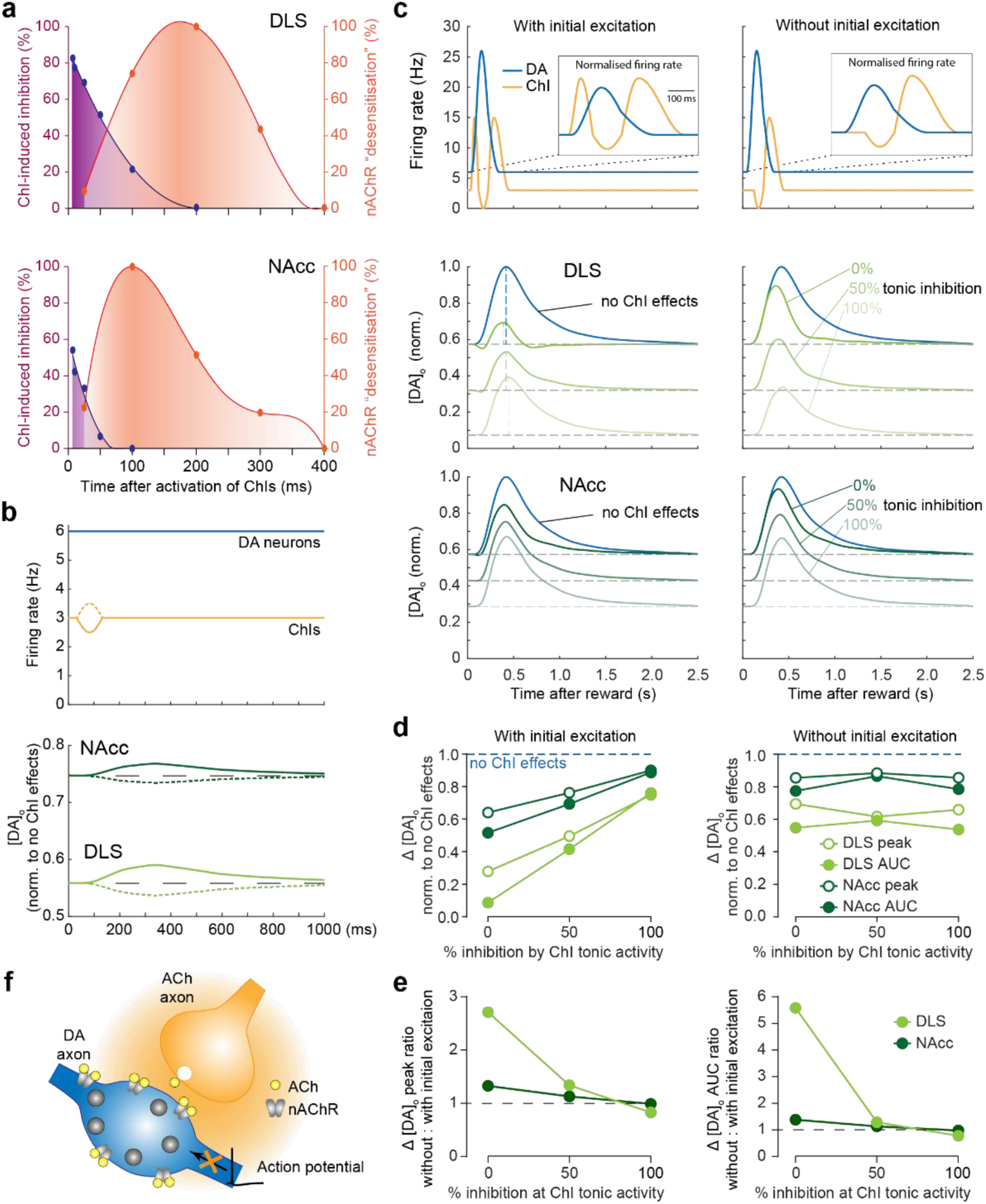
ChI-dependent attenuation of striatal DA release dominates in a computational model. **a,** Values used in the model for the strength of ChI-dependent depression of DA release (*purple*) and the normalised level of apparent nAChR desensitisation (orange) versus time after ChI activation in DLS (*upper*) and NAcc (*lower*). **b,** When firing rate of DA neurons (*blue*) is constant, a brief decrease (*solid*) or increase (*dotted*) of ChI activity (*yellow*) can respectively increase or decrease [DA]_o_ in DLS (*light green*) and NAcc (*dark green*). **c,** *Top row,* multiphasic ChI responses (*yellow*) with (*left*) or without (*right*) initial excitation phases, plus DA neuron burst activity (*blue*) from^22^ were inputted to predict striatal DA release (lower *rows*) in DLS and NAcc when the tonic level of ChI-dependent depression of DA release was set to 0% (*green*), 50% (*lighter green*), and 100% (lightest *green*). [DA]_o_ are normalised to those seen with zero tonic ChI effects (*blue*). **d,** Summary of peak [DA]_o_ (*open circle*) and area under [DA]_o_ curve (circle) in DLS (*light green*) and NAcc (*dark green*) with (*left*) and without (*right*) initial excitation in ChI multiphasic activity, when ChI-dependent attenuation was set to 0%, 50%, and 100% at tonic activity of ChIs. **e,** The ratio of peak [DA]_o_ (left) and area under [DA]_o_ curve (right) release when ChIs without and with initial excitation. **f,** Schematic showing that activation of ChIs and β2*-nAChRs on DA axons limits action potential propagation.

We first explored how changes to ChI tonic firing rate might modulate tonic [DA]_o_ in the absence of underlying changes in DA neuron firing rate. We set the tonic level of ChI-induced suppression of DA release to an arbitrary value of 50% of the maximum observed in each region, and normalised [DA]_o_ to the level seen without ChI effects. The model illustrates how transient (∼100 ms) multiphasic changes in ChI activity generate opposite changes in [DA]_o_ (**Fig. 6b**). This illustration helps to explain the observations *in vivo* that [DA]_o_ can be modified without an underlying change to DA neuron firing rate^6^. We then tested how multiphasic ChI activity modifies [DA]_o_ during concurrent burst activity in DA neurons^22^, both with and without the initial excitation that occurs in half of ChIs^21^ (**Fig. 6c**). We incorporated a range of levels of tonic ChI-induced depression of DA release (0, 50, 100%) prior to multiphasic activity which correspondingly reduced the baseline level of tonic DA release (**Fig. 6c**). In response to DA neuron burst activity, DA release was lowered by concurrent multiphasic activity in ChIs, and to a greater extent (i) when multiphasic activity included initial ChI excitation, and (ii) when baseline tonic ChI-induced depression of DA release was at a lower initial % strength against which the effect of multiphasic activity was offset, and (iii) in DLS than in NAc (**Fig. 6c-e**). The strongest predicted reduction in burst DA release arose particularly from the initial excitation in ChIs on a minimal prior background of tonic ChI activity, while in the absence of initial excitation, the strongest reduction in DA release resulted from depression of late DA release by ChI rebound activity. A nAChR desensitisation-like component played only a minor role in these outcomes, and in DLS more than NAc, and only when initial excitation was present not absent (**Extended Data Fig. 5**). Overall, the model suggests that multiphasic activity in ChIs attenuates DA release during phasic activity, particularly in DLS, and particularly in response to initial excitation.

The model also indicated that tonic ChI activity in the absence of multiphasic ChI activity, reflective of scenarios prior to learning^21,33^, reduces both tonic and burst-evoked levels of DA release (*blue lines,* **Extended Data Fig. 6a**). At low-intermediate levels of tonic suppression of DA release (<50%), multiphasic ChI activity further reduced phasic DA release. Only at extremely high levels of tonic suppression (50-100%), did ChI multiphasic activity enhance phasic DA release (**Extended Data Fig. 6a** and b), owing to relief of ChI-mediated depression of DA release enabled by a ChI pause. Therefore, the level of phasic striatal DA release will be a dynamic function of both tonic and multiphasic activity in ChIs.

## Discussion

Cholinergic interneurons have become a major focus of interest for their potential to regulate DA output. Here we reveal that activation of ChIs and nAChRs and initial depolarisation of the DA axon, at levels below or above those required to drive DA release, is followed by a strong refractory-like suppression of the subsequent excitability of DA axons, preventing further depolarisation by subsequent arriving activity, and limiting axonal Ca^2+^ summation and DA release. Our combined *ex vivo, in vivo* and *in silico* approaches together show that during physiological activity *in vivo*, ChIs then operate a strongly limiting effect that inversely scales DA release during activity in DA neurons. These findings support a major role for axonal integration as a mechanism to prevent neurotransmitter output, and also necessitate revision of a range of current prevailing views about how ACh governs DA output, in the following ways.

### Dynamic refractoriness of DA axons versus instantaneous depolarisation of DA axons

Activation of ChIs can rapidly depolarise DA axons via nAChR activation, action potential generation^8,34^ and drive instantaneous DA release^4,5^. However, little supporting evidence has been gained in very recent studies for the likelihood that ChIs drive DA release in vivo^35-37^, and our data set this against other outcomes of nAChR activation. We show that DA release is not the most feasible outcome of activation of ChIs/nAChRs *in vivo*, as it sits within a broader and opposing overall outcome of ongoing ACh-DA integration. We show that activity in ChIs can, after initial activation of nAChRs and axonal depolarisation, lead to an extended refractoriness of DA axons to subsequent activation by stimuli arriving up to ∼100 ms later. This could be mediated through either depolarisation-dependent ion channel inactivation or a form of shunting inhibition during or resulting from nAChR channel conductances. This striking refractoriness to further depolarization will provide an ongoing interruption of the relationship between spikes occurring in DA neurons and DA release to diminish the amplitude of DA output according to recent ChI activity.

We show that ChIs can more easily limit than activate DA release: ChIs can profoundly depress DA release following levels of ChI activation that are less than those required to drive detectable instantaneous DA release. Therefore, *in vivo*, when ChI activity is ongoing but also less synchronised than after artificial stimulations^4,5^, ChIs and ACh release are more likely to reach the lower levels that are sufficient to inhibit DA release than the higher levels required to trigger it. Indeed, *in vivo* we confirmed that the net outcome of activating nAChRs is to depress, and not drive, DA release. These interpretations do not exclude the possibility that activation of ChIs might also trigger DA release in certain scenarios *in vivo.* For example, if sufficient interval has lapsed since nAChR activation that the subsequent suppression of DA axon activity is minimised, and nAChRs are recovered from their activation-desensitisation cycle, and a population of ACh boutons or ChIs then fire in synchrony, a large instantaneous increase in ACh release and nAChR activation might result and be sufficient to trigger DA release in some microdomain. This might be more likely in NAc where the depression of DA release is weaker than DLS. Correspondingly, the relationships between concurrent ACh and DA signalling might show localised variation that can also vary over time.

Nonetheless, our data *in vivo* from detection of DA reporter GRAB-DA and axonal activity (using calcium imaging) show that activity in ChIs *in vivo* predominantly depresses DA release. These data, in conjunction with our modelling of DA release under different levels of activity in ChIs and DA neurons show that the impact of ChI activity on DA release will vary dynamically with tonic and multiphasic activity in ChIs, and will provide a continuously dynamic and inverse scaling factor on the amplitude of DA signals.

### Depression due to refractoriness versus depletion of vesicle pool

We identified that the limitation on subsequent DA release following nAChR activation was not due to a potential depletion of the DA vesicle pool that might follow an initial release event^9,10^. We saw that a low-level stimulation of ChIs that did not result in detectable DA release and could not have depleted the DA vesicle pool, nonetheless led to a profound depression of subsequent DA release, whether the subsequent stimulus was applied to ChIs or DA axons (see Figs 1-3). Therefore, the refractoriness on DA release after ChI activation is not via depletion of the DA vesicle pool.

### Refractoriness of subsequent DA release versus firing frequency-dependent filtering

The finding that nAChR activation limits subsequent DA axon depolarisation is not equivalent to the previous theory that ChIs provide a frequency filter on DA release in which the modulation of DA release by nAChRs varies with DA neuron firing frequency^2,38^. The interpretation of a filtering effect was based on the observation that during concurrent activation of DA axons and ChIs, nAChRs appeared to support DA release driven by trains of pulses at low but not high frequencies, akin to a form of ‘low-frequency-pass filter’. Our new observations now reveal that this apparent ‘frequency filtering’ is not in fact related to frequency of activity in DA neurons, but rather, will be an outcome of the refractoriness of DA axons to any stimuli occurring within ∼100 ms of nAChR activation. With concurrent local stimulation of DA axons and ChIs, DA release will be restricted to the initial summed DA_DA_+DA_ChI_, with little ensuing release for stimuli arriving up to ∼100 ms later, leading to minimal apparent sensitivity of DA release to subsequent DA neuron activity for frequencies ≥ 10 Hz. The apparent ‘low-frequency pass’ filtering effect was an inadvertent outcome of concurrently activating DA axons and ChIs. Rather, we show that activity in ChIs alone prevents subsequent DA release in a manner that depends on the interval between activity in ChIs and the ensuing activity in DA neurons, and not the frequency of firing of DA neurons *per se*. Since, *in vivo,* ChIs and DA neurons typically fire tonically at 3 – 10 Hz, and each DA axon is likely to be under the control of a network of overlapping ChI axons, DA release is likely under a predominant attenuation by ChIs and will scale inversely with recent ChI activity.

The frequency dependence of DA release *per se* will continue to be sculpted by other intrinsic mechanisms that regulate short-term facilitation, depression or summation of DA release, with other short-term depression mechanisms in particular persisting over much longer timescales (several seconds) than the mechanisms operated by nAChRs^11^.

### Attenuation versus enhancement of phasic DA release

Synchronised activation of a small population of ChIs by optogenetic or electrical activation or stimulation of their cortical or thalamic inputs has previously been shown to directly drive DA release^4,5,9,10,39^ leading to speculations that the initial excitation or rebound activity in multiphasic ChI activity acquired *in vivo* in response to reward or a reward-related cue during reinforcement learning will drive DA release. Our combined observations *ex vivo* and *in vivo* suggest that by contrast ChIs operate a predominantly limiting inverse scaling effect on DA release.

In addition, the pause in a ChI multiphasic response has previously been speculated to promote coincident burst-evoked DA, because high-frequency stimuli can induce more DA release when nAChRs are turned off^2,40^. However, while a reduction in ChI activity from tonic levels or after an initial phase of excitation in a multiphasic burst may relieve the inhibition on DA release, its attenuation for up to 100 ms will persist into a short pause phase and continue to place some limitations on DA release. This effect will decay during the pause and scale with prior ChI activity, and will be more profound with greater initial excitation. Our model also shows that while ChIs inversely regulate the amplitude or scale of DA signals it does not modify their kinetics.

### Regional heterogeneity of ChI-induced depression of DA release

ChIs prevent DA axonal depolarisation more strongly and for longer durations in DLS than NAcc which will lead to critical distinctions in function. DA neurons encode reward predictions and their errors through phasic firing frequency but despite largely similar events in SNc^1^ and VTA^41^ during learning, there are larger amplitude DA release events detected in ventral (from VTA) than dorsal (from SNc) striatum *in* vivo^42,43^. This discrepancy does not occur *ex vivo,* where local striatal stimulations with single pulse stimuli typically evoke lower [DA]_o_ in NAc than DLS^17,44^, indicating that a dynamic circuit(s) found *in vivo* is more permissive for phasic DA release in NAc than DLS. The lesser refractoriness of DA release after ChI/nAChR activity in NAc than in DLS is a candidate explanation (**Fig 6b,c**), and in turn suggests that ChI-induced depression of DA release may play a critical role in regulating how reward prediction errors translate differently to DA release in different striatal regions^43,45,46^ and thus contribute to striatal learning^47^.

In summary, the extended apparent refractoriness of DA axons after nAChR-mediated depolarisation is a major mechanism through which striatal β2-nAChRs attenuate DA axon excitability to limit the level of DA release. This mechanism will operate dynamically on DA axons according to ChI activity to dominate as a gain mechanism that inversely scales DA output continuously and dynamically according to the recent history of ChI activity.

## Materials and Methods

### Animals

Mice used in *ex vivo* experiments and *in vivo DA* recordings were adult male C57Bl6/J mice (Charles River, UK) (21-40 days), heterozygous ChAT-Cre:Ai32 (6-16 weeks), heterozygous DAT-IRES-Cre (8-16 weeks), heterozygous DAT-Cre:Ai95D (4-7 weeks), or heterozygous DAT-Cre:ChAT-Cre mice (8-12 weeks). ChAT-Cre^+/+^ mice (B6;129S6-Chat^tm2(cre)Lowl^/J, JAX stock number 006410) were crossed with Ai32^+/+^ mice (B6;129S-Gt(ROSA) 26Sor^tm32(CAG-COP4*H134R/EYFP)Hze/^J, JAX stock number 012569) to produce heterozygote ChAT-Cre:Ai32 mice. DAT-IRES-Cre^+/-^ mice (B6.SJL-Slc6^a3tm1.1(cre)Bkmn^/J, JAX stock number 006660) were injected in midbrain with pAAV-double floxed-hChR2(H134R)-EYFP-WPRE-pA for ChR2 expression in DA axons (8-16 weeks). DAT-IRES-Cre^+/-^ mice were crossed with Ai95D^+/+^ mice (B6:129S-Gt(ROSA)26Sor^tm95.1(CAG-GCaMP6f)Hze^/J) to create heterozygote DAT-Cre:Ai95D mice. Male C57BL/6N mice (Charles River, Beijing, China) (42-50 days) were used for behavioural experiments.

Animals were group-housed and maintained on a 12-hour light/dark cycle with *ad libitum* access to food and water. The procedures for *ex vivo* recordings and anaesthetised *in vivo* DA recordings were performed in accordance with Animals (Scientific Procedures) Act 1986 (Amended 2012) with ethical approval from the University of Oxford, and under authority of a Project Licence granted by the UK Home Office. Behavioural experiments were performed using protocols approved by the Animal Care & Use Committees at the Chinese Institute for Brain Research (#CIBR-IACUC-007) and were performed in accordance with the guidelines established by US National Institutes of Health.

### Virus injection

Mice were placed in a stereotaxic frame under isoflurane anaesthesia and a craniotomy was made above the injection site. Injections of 1 μL virus were given either unilaterally or bilaterally in either VTA (co-ordinates from Bregma in mm: AP −3.1, ML ± 0.5, DV −4.4) or in the SNc (from Bregma in mm: AP −3.5, ML ± 1.2, DV −4.0) using a 2.5 μL 33-gauge Hamilton syringe at 0.2 µL/min with a microinjector. The syringe was left in place for 5 min following each injection, then retracted slowly. Animals were maintained for at least 3 weeks following surgery to allow virus expression in striatum.

For expression of ChR2 or ASAP3 in DA axons, heterozygote DAT-IRES-Cre mice were injected intracerebrally with either a Cre-inducible recombinant AAV serotype 5 vector containing an inverted gene for channelrhodopsin-2 fused in-frame with a gene encoding enhanced yellow fluorescent protein (pAAV-double floxed-hChR2(H134R)-EYFP-WPRE-pA) (titre = 1E+12 vg/ml, University of North Carolina Vector Core), or a Cre-inducible recombinant AAV serotype 5 vector containing a EF1A promoter and an inverted gene for ASAP3 (titre = 2.4E+12 vg/ml) without a soma-targeting signal and a WPRE RNA-stabilizing element (Stanford Gene Vector and Virus Core).

For dual optogenetic experiments, heterozygote DAT-Cre:ChAT-Cre mice were injected intracerebrally with a Cre-inducible recombinant AAV serotype 5 vector containing an inverted gene for Chrimson (ssAAV-5/2-hEF1α/hTLV1-dlox-ChrimsonR_tdTomato(rev)-dlox-WPRE-bGHp(A), v288-5 ETH Zurich) into midbrain and a Cre-inducible recombinant AAV serotype 2 vector containing an inverted gene for channelrhodopsin-2 fused in-frame with a gene encoding enhanced yellow fluorescent protein (pAAV-double floxed-hChR2(H134R)-EYFP-WPRE-pA) (titre = 1E+12 vg/ml, University of North Carolina Vector Core) into the striatum.

For expression of GRAB-DA_2m_, wild-type C57BL/6J mice were injected intracerebrally with AAV serotype 5 vector containing GRAB-DA_2m_ (titre = 1E+13 vg/mL, BrainVTA, China) into dorsal striatum.

### Ex Vivo Slice Voltammetry and Stimulation

For fast-scan cyclic voltammetry (FCV) in acute coronal slices, animals were anaesthetised with isoflurane. Brains were quickly removed into ice-cold, high Mg^2+^ cutting solution containing in mM: 85 NaCl, 25 NaHCO_3_, 2.5 KCl, 1.25 NaH_2_PO4, 0.5 CaCl2, 7 MgCl_2_, 10 glucose, 65 sucrose. Brains were then blocked, and 300 µm coronal slices were cut on a vibratome (Leica VT1200S) between +1.5 to +0.5 mm from bregma containing caudate-putamen and nucleus accumbens. Slices recovered at 32°C for 30-40 min after dissection and were subsequently kept at room temperature. Slices were maintained and recorded in artificial cerebrospinal fluid (aCSF) containing in mM: 130 NaCl, 25 NaHCO_3_, 2.5 KCl, 1.25 NaH_2_PO_4_, 2.5 CaCl_2_, 2 MgCl_2_, 10 glucose. The aCSF was saturated with 95% O_2_/ 5% CO_2_; recordings were made at 32-33 °C. Extracellular DA concentration ([DA]_o_) was measured using FCV with 7 µm-diameter carbon fiber microelectrodes (CFMs; tip length 50-100 µm) and a Millar voltammeter (Julian Millar, Barts and the London School of Medicine and Dentistry) as previously^48^. The voltage was applied as a triangular waveform (- 0.7 to +1.3 V range versus Ag/AgCl) at a scan rate of 800 V/s, and data were sampled at 8 Hz.

For optogenetic stimulations, ChR2-expressing ChIs or DA axons were activated using a 473 nm diode laser (DL-473, Rapp Optoelectronic) coupled to the microscope with a fiber optic cable (200 µm multimode, NA 0.22). Spot illumination had a 30 µm diameter under x40 immersion objective. Laser pulses were 2 ms duration, 5-23 mW/mm^2^ at specimen. Chrimson-expressing DA axons were activated using a LED with a 585 ± 22 nm filter. LED pulses were 2 ms duration, 2.5-3.4 mW/mm^2^. The Lstim_0_ was achieved by lowering the laser intensity to the point at which there was no detectable evoked FCV signal above noise, in on-line or off-line analyses (Extended Fig 1).

For electrical stimulations, 0.65 mA current (200 µs width) was delivered through a surface bipolar concentric Pt/Ir electrode (125 µm outer, 25 µm inner diameter, FHC, USA) placed ∼100 µm from the recording electrode. The Estim_50_ was the stimulation current at which evoked [DA]_o_ was ∼50% of that seen with normal stimulation (0.65 mA) during on-line analysis. Stimulations were timed to avoid FCV scans.

### Calcium Imaging Ex Vivo

As in our previous study^11,14^, an Olympus BX51Wl microscope equipped with a OptoLED Lite system (CAIRN Research), Prime Scientific CMOS (sCMOS) Camera (Teledyne Photometrics), and a x40/0.8 NA water-objective (Olympus UK) was used for wide-field fluorescence imaging of GCaMP6f in dopaminergic axons in DLS in *ex vivo* slices in response to single and trains (4 pulses, 100 Hz) of electrical stimulus pulses. Images were acquired at 16.6 Hz frame rate every 2.5 min using Micro-Manager 1.4, with stimulation and recording synchronised using custom-written procedures in Igor Pro 6 (WaveMetrics) and an ITC-18 A/D board (Instrutech). Image files were analysed with Matlab R2019b and Fiji 1.5. We extracted fluorescence intensity from the region of interest (ROI) of 25 µm * 25 µm which was 50 µm away from the electrical stimulating electrode tip. Ca^2+^ transients were bleach-corrected by fitting an exponential curve function through both the baseline (2 s prior to stimulation) and the last 1 s in a 7.2 s recording window. The order of single and train stimulations was alternated and equally distributed and data were collected in duplicate before and after a change in extracellular experimental condition. Data are expressed as ΔF/F where F is the fitted curve.

### Voltage Sensor Imaging Ex Vivo

An Olympus BX51Wl microscope equipped with a OptoLED Lite system (CAIRN Research), an iXon EMCCD Camera (ANDOR), and a x40/0.8 NA water-objective (Olympus UK) was used for wide-field flEMCCD Camera (ANDOR), and a x40/0.8 NA water-objective (Olym*ex vivo* slices in response to single and trains (4 pulses, 500 Hz) of electrical stimulus pulses. Images were acquired at 660 Hz frame rate every 2.5 min using Micro-Manager 1.4, with stimulation and recording synchronised using PClamp. Image files were analysed with Matlab R2019b and Fiji 1.5. We extracted fluorescence intensity from the region of interest (∼5 µm * 5 µm). The ASPA3 transients were bleach-corrected by fitting an exponential curve function. The order of single and train stimulations was alternated and equally distributed and data were collected in duplicate before and after a change in extracellular experimental condition. Data are expressed as ΔF/F where F is the fitted curve.

### Anesthetised In Vivo Recordings

Wild type mice were anesthetised with urethane (1.4–1.9 g/kg, i.p.; Biolab), supplemented with additional urethane (0.2 g/kg) every 1-2 hr as required. All wounds and pressure points were infiltrated with bupivacaine (0.5%). Upon reaching surgical anaesthesia, the head was fixed in a stereotaxic frame (Kopf, USA). Core temperature was maintained at 35-36 °C using a homeothermic blanket and monitored via a rectal probe (TR-100, Fine Science Tools). Mecamylamine (2 mg/kg) was injected *i.p.* to block nAChRs in the striatum.

### *In Vivo* Voltammetry

A round piece of skull overlying the left hemisphere was removed to target the DLS (AP +1.0 mm, ML 1.6 mm, DV 2.2 mm to Bregma). A stimulating and recording array consisting of a carbon-fiber microelectrode and a bipolar stimulating electrode (MS303/3-A/SPC, P1 Technology) was positioned in the DLS. The Ag/AgCl reference electrode was implanted in another part of the forebrain. [DA]_o_ was measured using FCV with 7 µm-diameter carbon fiber microelectrodes (CFMs; tip length 50-100 µm) and a Tarheel system (University of Washington, Seattle, US). The voltage was applied as a triangular waveform (-0.4 to +1.3 V range versus Ag/AgCl) at a scan rate of 400 V/s and data were sampled at 10 Hz. The location of the tip of FCV electrode was confirmed histologically. For striatal electrical stimulation, 0.65 mA current (200 µs) was delivered through a bipolar stimulating electrode (0.005 inch, MS303/3-A/SPC, P1 Technologies). The stimulating electrode tips were separated by ∼500 µm and were glued to the FCV recording electrode to fix the tip of the FCV electrode between the two stimulating poles.

### *In vivo* Fiber Photometry

Round pieces of skull overlying the left hemisphere were removed to allow access to the DLS (AP +1.0 mm, ML 1.6 mm, DV 2.2 mm to Bregma) and SNc (AP -3.1mm, ML 0.8mm, DV 4.3mm to Bregma). The injection and recording array, consisting of a glass pipette and a 200 µm diameter fibre, was positioned in the DLS. GCaMP6f expressed in DA axons or GRAB-DA_2m_ was activated with 480 nm light (76 µW), and the intensity of GCaMP6f or GRAB-DA_2m_ emission was sampled at 40 Hz with Neurophotometrics (FP3001).

For midbrain electrical stimulation, 0.5 mA current (500 µs) was delivered through a bipolar stimulating electrode (0.005 inch, MS303/3-A/SPC, P1 Technologies) at 0.1 Hz. The stimulating electrode tips were separated by ∼500 µm.

### Behavioural Recordings

#### Cannulae Placements

Male C57BL/6N mice (42-50 days) were anesthetised with isoflurane (5% induction; 1.5 - 2% maintenance) and placed on a stereotaxic frame for surgery. Bilateral injection needles (O.D. 0.21 mm, I.D. 0.11 mm, RWD, China) with the guide cannula (O.D. 0.41 mm, I.D. 0.25 mm, RWD, China) were implanted to the dorsal striatum either vertically or at a small angle from the vertical, with the tip of each cannula aimed at coordinates: AP +1.0 mm; ML +/- 1.6 mm to bregma; DV -2.4 mm (from dura). Mice recovered for three days after surgery.

#### Conditioned Place Preference Testing

In CPP experiments, mice were placed in a 40 cm × 40 cm transparent plexiglass arena which was divided into two equal chambers separated by doorway. The chambers were decorated with either horizontal or vertical stripes. The movement of animals was recorded and analysed with Smart V3.0 tracking software (Panlab, Spain). On day 1, mice were allowed to freely shuttle between two chambers to assess place preference at baseline, expressed as % time spent in right chamber. The mice were conditioned on days 2 and 3, when animals received alternating bilateral striatal injection with either mecamylamine (10 µg/side) or saline vehicle (0.9%) in a volume of 0.5 µl over 1 min in AM and PM. Animals were then constrained respectively in the right or left chamber for 20 min. The treatments were counterbalanced for time of day. On day 4, the post-conditioning chamber preference was calculated as the % of time spent in the right mecamylamine-associated chamber compared to on pre-conditioning day 1. For the next two days (days 5-6), animals received bilateral saline injection and explored both chambers for 20 min after which place preference was extinguished. The conditioning procedure was then repeated but for bilateral saline for both chambers, with a pre-conditioning test on day 7, two days of conditioning on days 8-9, and a post-conditioning test on day 10. To minimise place preference bias at baseline, the five animals in each test showing least place preference on the pre-conditioning day (mecamylamine 42%-58%; control 45%-55%) were selected for subsequent conditioning. For open field experiments, mice received bilateral striatal injection of either saline vehicle or mecamylamine (10 µg/side), and were placed into the open field chamber to assess total running distance and average velocity within 20 minutes.

### Immunocytochemistry

After voltammetry recordings in acute slices, slices were fixed in 4% paraformaldehyde dissolved in PBS containing 0.2% picric acid. Slices were fixed overnight at 4°C and then stored in PBS. Free-floating sections were then washed x5 in PBS for 5 min and incubated in 0.5% Triton X-100 and 10% normal donkey serum.

ChIs expressing ChR2-eYFP were identified as ChAT-immunoreactive as previously (Zhang et al., 2018). Fixed and rinsed slices were incubated overnight with goat anti-ChAT 1:100 antibody (Millipore) or for five days with goat anti-ChAT 1:200 (AMCA, Jackson Immuno Research Europe) dissolved in PBS containing 0.5% Triton X-100 and 3% normal donkey serum. Sections were then washed x5 with PBS for 5 min and incubated for 2 hr at room temperature with 1:1000 Alexa Fluor 568 donkey anti-goat antibody (Invitrogen) dissolved in PBS containing 0.5% Triton X-100 and 3% normal donkey serum.

DA neurons co-expressing ChR2-eYFP, GCaMP6f-eYFP, ASAP3 or Chrimson and striatal DA axons were identified by immunoreactivity to tyrosine hydroxylase (TH) as previously^11,14^. Fixed and rinsed slices were incubated overnight with 1:2000 rabbit anti-TH antibody (Sigma) dissolved in PBS containing 0.5% Triton X-100, 1% normal goat serum and 1% foetal bovine serum. Sections were then washed x5 with PBS for 5 min and incubated for 2 h at room temperature with 1:1000 (DyLight 594 goat anti-rabbit antibody, Jackson, or CoraLite488 goat anti-rabbit antibody, Proteintech) dissolved in PBS containing 0.5% Triton X-100, 1% normal goat serum and 1% foetal bovine serum.

Sections processed as above were then washed with PBS and mounted on gelled slides with Vectashield mounting medium (Vector Labs) and imaged at 20x, N.A. 0.8, using a Zeiss LSM880 confocal microscope system running ZEN black version 2.3 (Zeiss), or on a confocal microscope system (FV1000 IX81; Olympus) using a 20×/0.75 NA objective and Fluoview software (Olympus). Maximum intensity projection from a z-stack of height 30 µm was captured individually and the stack of the pictures were compressed. Red fluorescence (TH and ChAT) was captured at 638–759 nm with 633 nm excitation. Green fluorescence (GCaMP, ChR2, and ASAP3) was captured at 493–630 nm with 488 nm excitation.

To verify carbon-fibre locations in dorsal striatum for *in vivo* FCV recordings, anaesthetised mice were sacrificed and brains were quickly removed and fixed in 4% paraformaldehyde (PFA) overnight. The fixed brains were then sectioned into 50 μm slices using a Vibratome (Leica). Slices were rinsed with PBS 3 times and mounted on glass slides and then imaged under microscope to identify the location of recording sites.

To verify placements of intrastriatal injection cannulae in behavioural experiments, mice were anesthetised with i.p. injection of Avertin (250 mg/kg body weight) and transcardially perfused with saline and 4% PFA. Brains were dissected and post-fixed overnight in 4% PFA then dehydrated by 30% sucrose for 24 hours. The fixed brains were then frozen-sectioned into 50 μm slices using a Vibratome (Leica). To verify the placement of cannulae, slices were stained with immunoreactivity to TH as slices from *ex vivo* experiments and mounted on glass slides. The slices were imaged under inverted confocal microscope (Zeiss) with 405 nm laser for excitation.

### Drugs

DHβE and mecamylamine hydrochIoride for *ex vivo* and anaesthetised *in vivo* experiments were purchased from Tocris Bioscience (UK). Mecamylamine hydrochIoride for behavioural experiments were purchased from Sigma Aldrich (China). All other chemicals were purchased from Sigma Aldrich (UK). Pharmacological drugs for *ex vivo* experiments were prepared in distilled deionised water as stock aliquots at 1000x final concentrations and stored at -20 ⁰C. Drug stocks were then diluted to the final concentration in carbogenated aCSF immediately before use and were bath-applied. Drugs for *in vivo* experiments were dissolved in sterilised saline to final concertation.

### Computational Model

The computational model was written in MATLAB and is available at https://github.com/Yanfeng-Zhang/ChI_prevent_DA. The model included: (1) the dynamic strength of ChI-dependent depression, determined from the ratio of [DA]_o_ evoked at a second stimulus before and after antagonizing β2*-nAChRs with DHβE (from Fig. 3d,h; DAT-Cre, light stimulus) fitted to a polynomial curve. DLS: *y* = 1.97e + 5*x*^2^ - 0.00833*x* + 0.872, R^2^ = 0.99; NAcc: *y* = 8.71e + 5*x*^2^ - 0.0149*x* + 0.611, R^2^ = 0.96; (2) the profile of apparent nAChR “desensitisation” (Fig. 4j,l) estimated from the change in the difference between [DA]_o_ evoked by 3 pulse and 1 pulse at a second stimulus (from Fig. 4i,k), normalised to a maximum and fitted with polynomial curves. DLS: y = 2e- 05*x*^2^ – 0.0087*x* +0.88 NAcc: y = -9e-10*x^4^* + 9e-07*x^3^* - 0.0003*x*^2^ + 0.039*x* - 0.5654; and (3) the dynamic release and uptake profile of [DA]_o_ seen after a single electrical stimulation *ex vivo* in order to model [DA]_o_ *in vivo* as a scalar product with DA neuron activity. We also included in the model a range of potential levels of background tonic ChI-dependent suppression of DA release (0, 50, 100%) arising from tonic activity in ChIs.

We excluded a potential component of DA release that can be driven by synchronised activation of ChIs in some experimental scenarios (Fig. 1)^4,5^, since we found that the threshold for nAChR-mediated suppression of DA release is lower than that required to drive release *ex vivo* (see Fig. 1) and was met *in vivo* after discrete striatal stimulation (see Fig. 4). We also excluded short-term depression arising from nAChR-independent mechanisms, as this co-varies only minimally on the short and rapid timescales relevant to the timescale of multiphasic activities^11^.

### Quantification and Statistical Analysis

Statistical analyses used GraphPad Prism 6.0. Data are expressed as mean ± standard error of the mean (SEM). The N value is the number of animals, *n* value is the number of individual recordings. One-sample t test, t-test, and two-way ANOVA were used.

### Data availability

The datasets generated during and/or analysed during the current study are available.

## Acknowledgments

We thank Alan Wainman from the Dunn School Bioimaging facility and Dr Dave Bergin, Ross McLeod, Dr Jeffery Stedehouder, and Shinil Raina for assisting with immunohistochemistry imaging. We thank Neuroscience Gene Vector and Virus Core (GVVC) at Stanford University (USA) for viral packaging of ASAP3. The work was supported by Parkinson’s UK (J-1403; G-1305), the Medical Research Council (MR/K013866/1, MR/J004324/1), and Aligning Science Across Parkinson’s (ASAP-020370).

## Author contributions

Y.-F.Z. conceived the study. Y.-F.Z. and S.J.C. designed the experiments. Y.-F.Z. performed experiments, collected and analysed the data, with contributions from Q.Q. and Y.H., and built the computational model. Y.-F.Z., S.J.C. and M.J. designed behavioural experiments, P.L. and M.J. performed the behavioural experiments and collected and analysed the data. P.Z. and A.L. provided expertise and assistance with fibre photometry recordings. G.Z. and M.Z.L. constructed and tested the ASAP3 AAV. Y.-F.Z. and E.O.M. performed voltage sensor imaging. Y.-F.Z. and S.J.C supervised the study. Y.-F.Z. and S.J.C prepared the manuscript with input from the other authors.

## Competing interests

Y.-F.Z. is the owner of patent (WO2023175357A1).

---

**Extended Data Fig 1.**
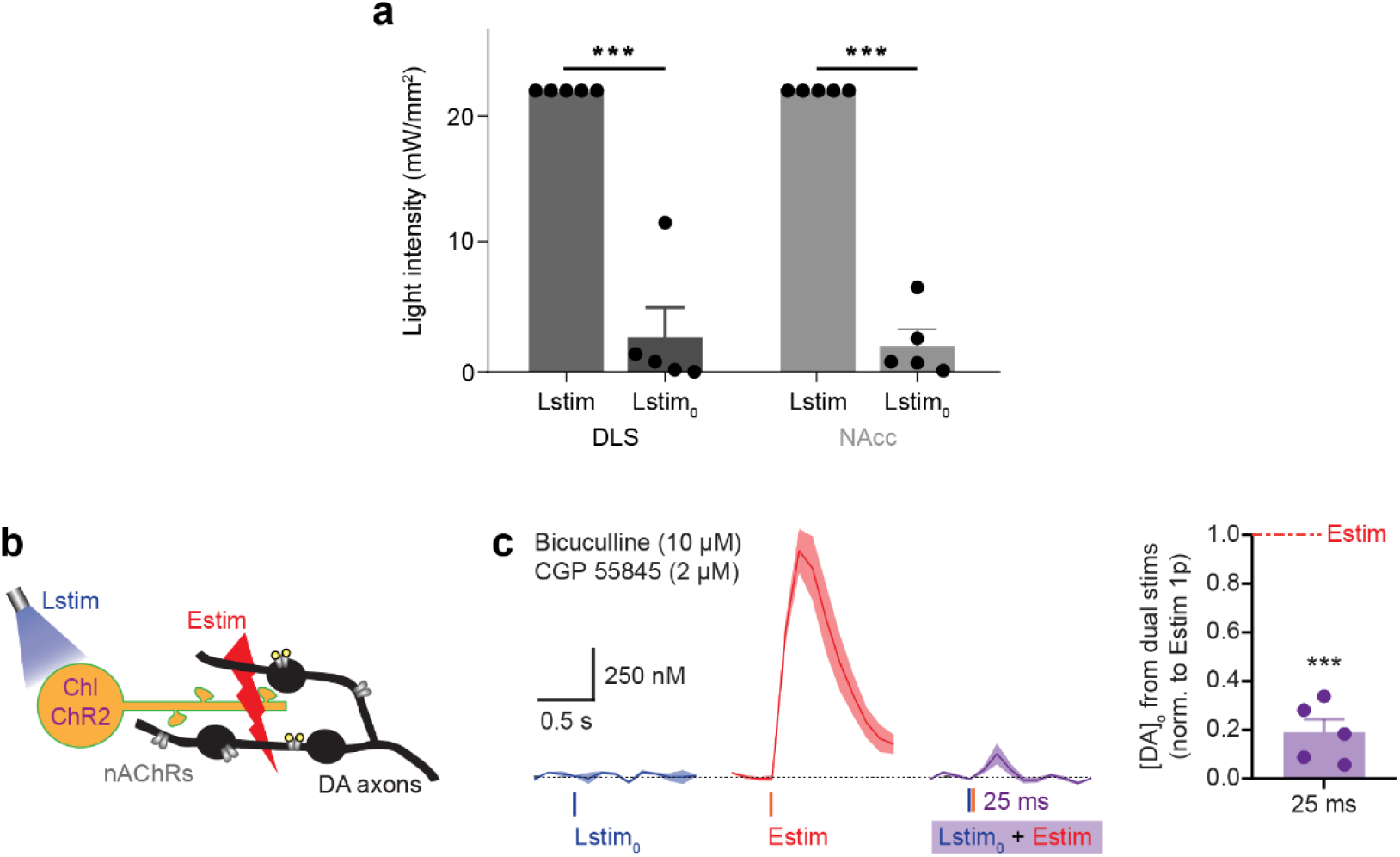
Light stimulation (Lstim_0_) intensities that are sub-threshold for evoking detectable DA release. **a,** The intensities of Lstim_0_ used to stimulate ChIs in ChAT-Cre:Ai32 mice in Figure 1 were significantly lower than Lstim in both DLS (black) and NAcc (grey). N = 5 animals. ***P<0.001, one sample t-test. **b**, Schematic of stimulation configuration for data in c. Blue light stimulation (Lstim) of ChR2-eYFP-expressing ChIs, subsequent local electrical stimulation (Estim) in striatal slices from ChAT-Cre:Ai32 mice. **c**, Mean transients (± SEM) from representative experiments in DLS showing [DA]_o_ evoked by either a subthreshold light pulse (Lstim_0_, blue), a single Estim (red), or paired Lstim and Estim (purple) at an ISI of 25 ms when GABA_A_ and GABA_B_ receptors were inhibited using bicuculline (10 µM) and CGP 55845 hydrochloride (2 µM) respectively. Mean peak [DA]_o_ (± SEM) evoked by the dual stimuli normalised to [DA]_o_ evoked by a single Estim (n = 5 recordings, N = 3 animals) when GABA receptors are inhibited. ***P<0.001, One sample t-test versus Estim.

**Extended Data Fig 2.**
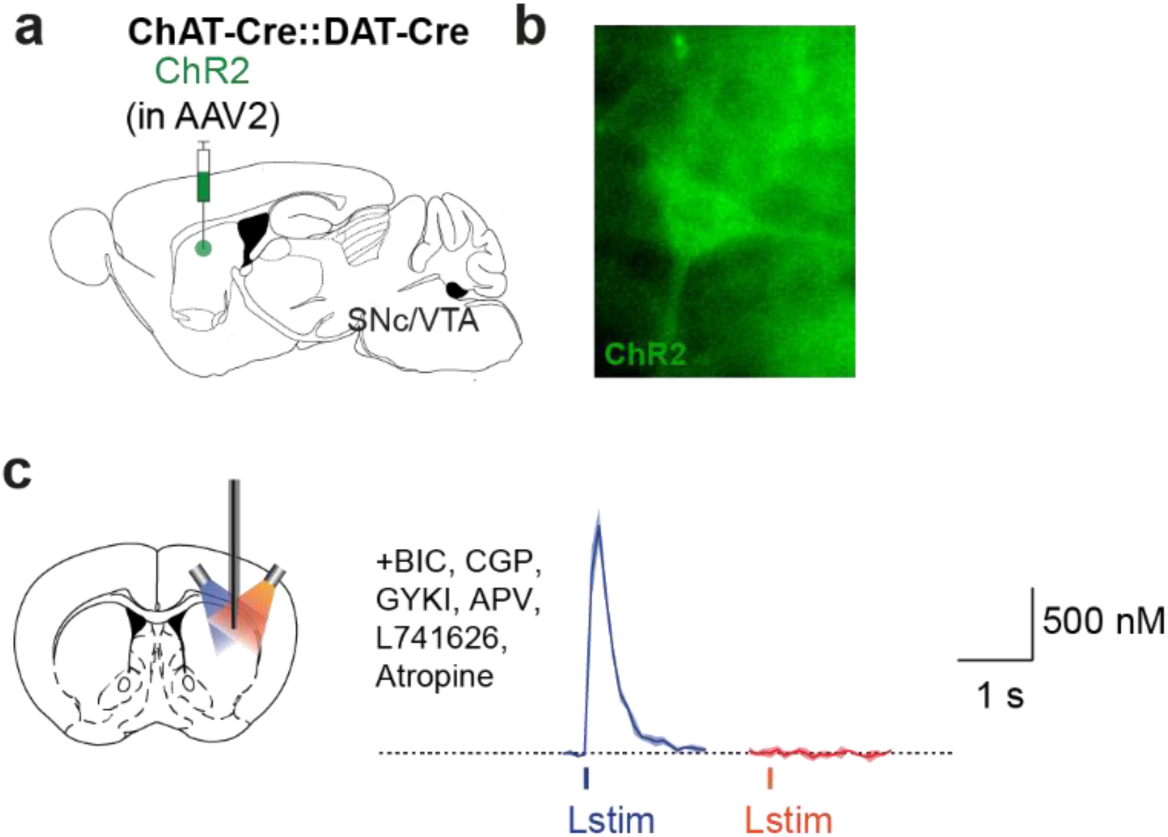
Orange LED has minimum ability to trigger DA release via ChR2-expressing Chls. **a**, Schematic of virus injection. **b**, A Chl expresses ChR2. **C**, *Left*, Schematic of stimulating configuration. *Right*, mean transients from representative experiments of [DA]_o_ (± SEM) evoked by a single pulse of blue Lstim (*blue lines*), or orange Lstim (orange *lines*). Blue light activated ChR2-eYFP-expressing ChIs to drive DA release. Orange light did not drive observable DA release via Chls. Antagonists of GABA_A_, GABA_B_, AMPA, NMDA, D_2_, and mAChRs were present.

**Extended Data Fig 3.**
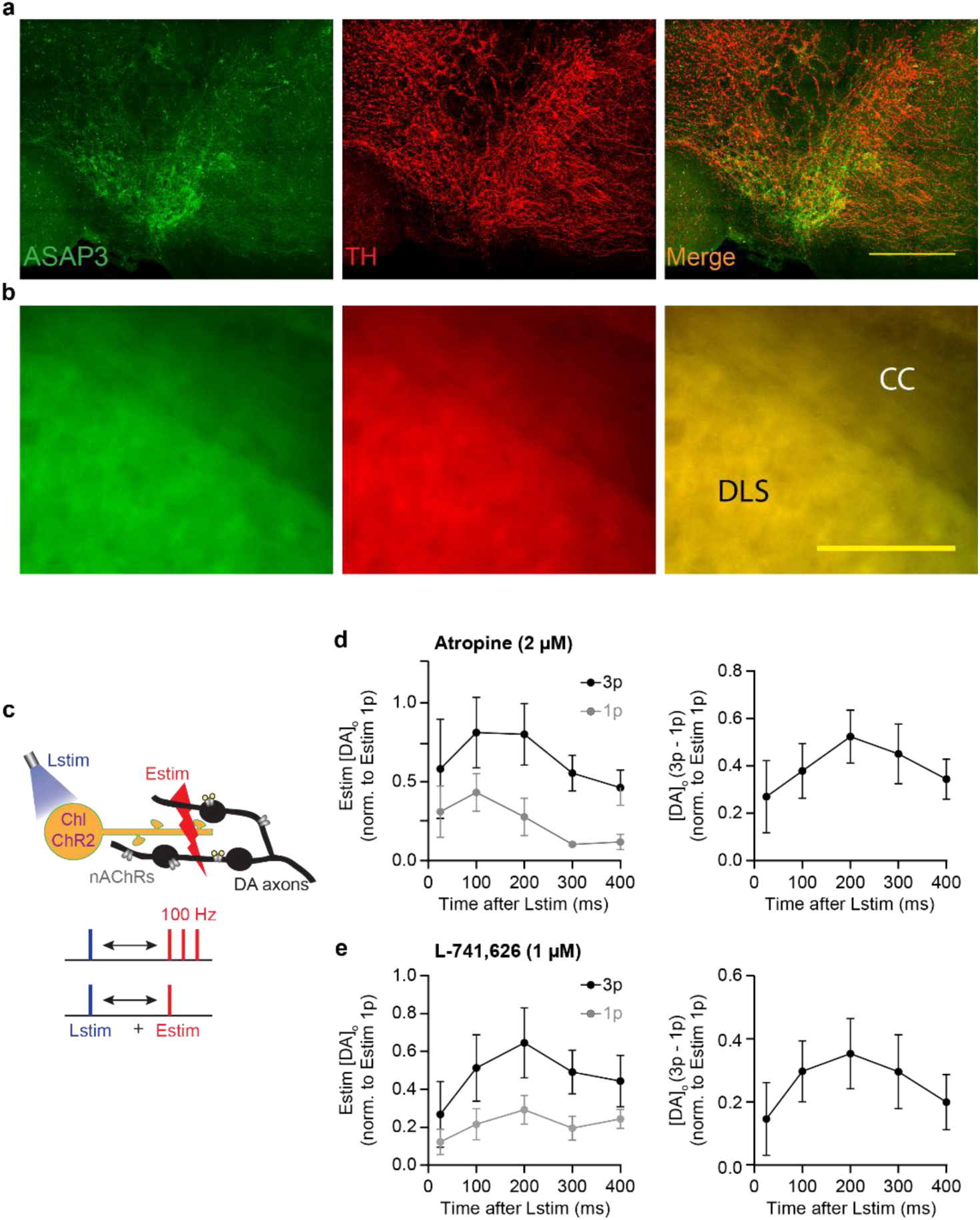
Expression of ASAP3 in DA neurons and the dynamic DA release response after light activation of ChIs (100-400 ms interval) persists in DLS in the presence of antagonists for either muscarinic receptors or D_2_ receptors. **a**, Expression of ASAP3-GFP after viral injection in DA neurons within midbrain co-labelled for TH-immunoreactivity. **b**, The co-expression of ASAP3 and TH in dopamine axons in the DLS. CC: corpus callosum (scale bar, 400 µm). **c**, Schematic of stimulation configuration used in d and e. Blue light stimulation (*Lstim*) of ChR2-eYFP-expressing ChIs, subsequent local electrical stimulation (*Estim*) in striatal slices from ChAT-Cre:Ai32 mice. **d,e**, *Left,* mean peak [DA]_o_ (± SEM) evoked in DLS after a Lstim by a subsequent Estim of either 1p (*grey*) or 3p 100 Hz (*black*) at a range of ISI, normalised to [DA]_o_ evoked by a single Estim, after antagonising either muscarinic receptors with atropine (2 µM) (**d**, n = 5 recordings, N = 3 animals) or D_2_ receptors with L-741,626 (1 µM) (**e**, n = 5 recordings, N = 4 animals). *Right*, differences between [DA]_o_ evoked by 3p and 1p Estims. Significant effect of IPI seen in bell-shaped curves for both, P<0.001, One-way ANOVA.

**Extended Data Fig 4.**
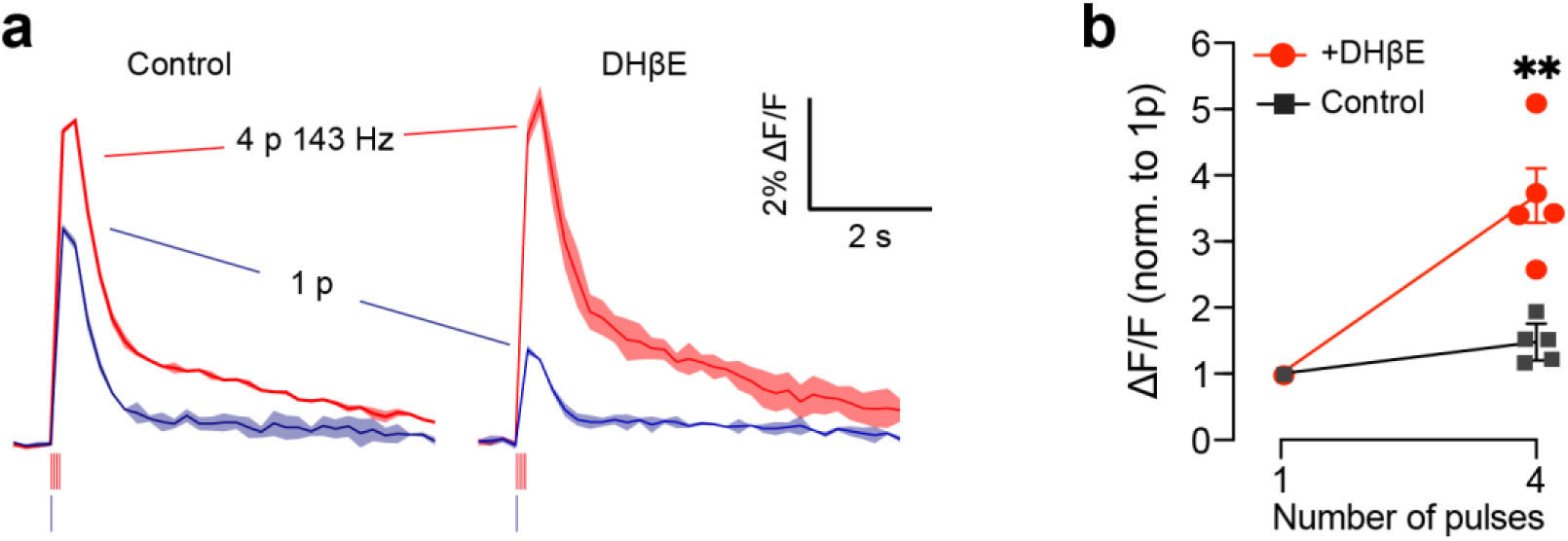
Activating nAChRs *ex vivo* prevents calcium entry into DA axons from subsequent electrical stimulation. **a,** Examples of calcium imaging responses (changes to GCaMP6f fluorescence, ΔF/F) (mean ± SEM from duplicates) in a DA axon population imaged in striatum in response to single or trains of 4 electrical pulses (4p) at 143 Hz (7 ms inter-stimulation interval) in control conditions (left) and in the presence of DHβE (1 μM) (right). **b,** Mean peak values (± SEM) for GCaMP6f ΔF/F vs. pulse numbers. Data are normalised to value for 1 pulse (n = 5 recordings, N = 2 animals). ** P < 0.01 paired t-test [DA]_o_ from 4 pulses of stimulation.

**Extended Data Fig 5.**
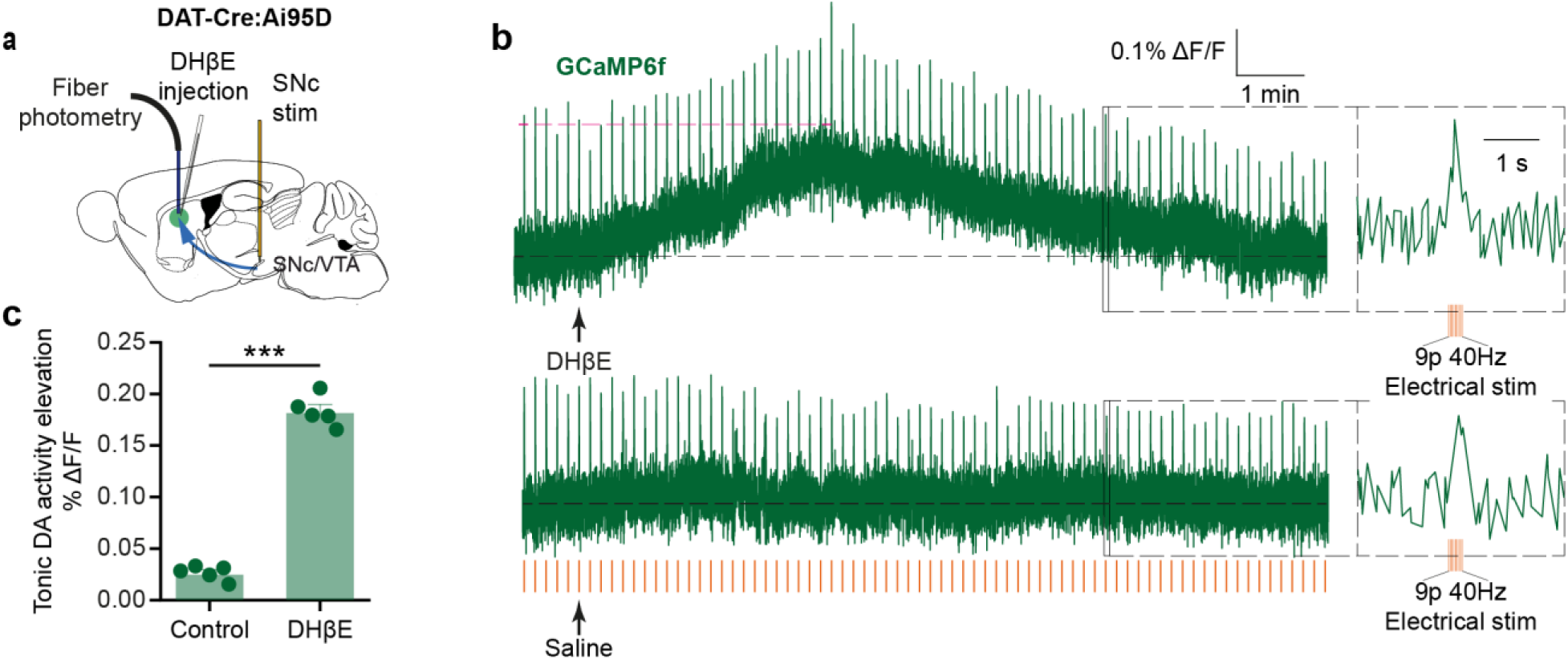
Striatal antagonism of nAChRs enhances calcium influx into DA axons *in vivo*. **a,** Schematic of configuration of fibre photometry recording in DLS and stimulation in SNc. **b**, Example traces of fibre photometry recording of GCaMP6f signal with local injection of DHβE (0.07µg in 200 nl saline) or saline (200 nl) showing tonic elevation after DHβE. Intermittent electrical stimulation (9 pulses at 40 Hz, red lines) was applied at 0.1 Hz in SNc. **c**, Mean peak change in tonic DA axon activity (± SEM) after local injection of DHβE or saline. ***P < 0.001 paired t-test (n = 5 from N = 3 animals).

**Extended Data Fig 6.**
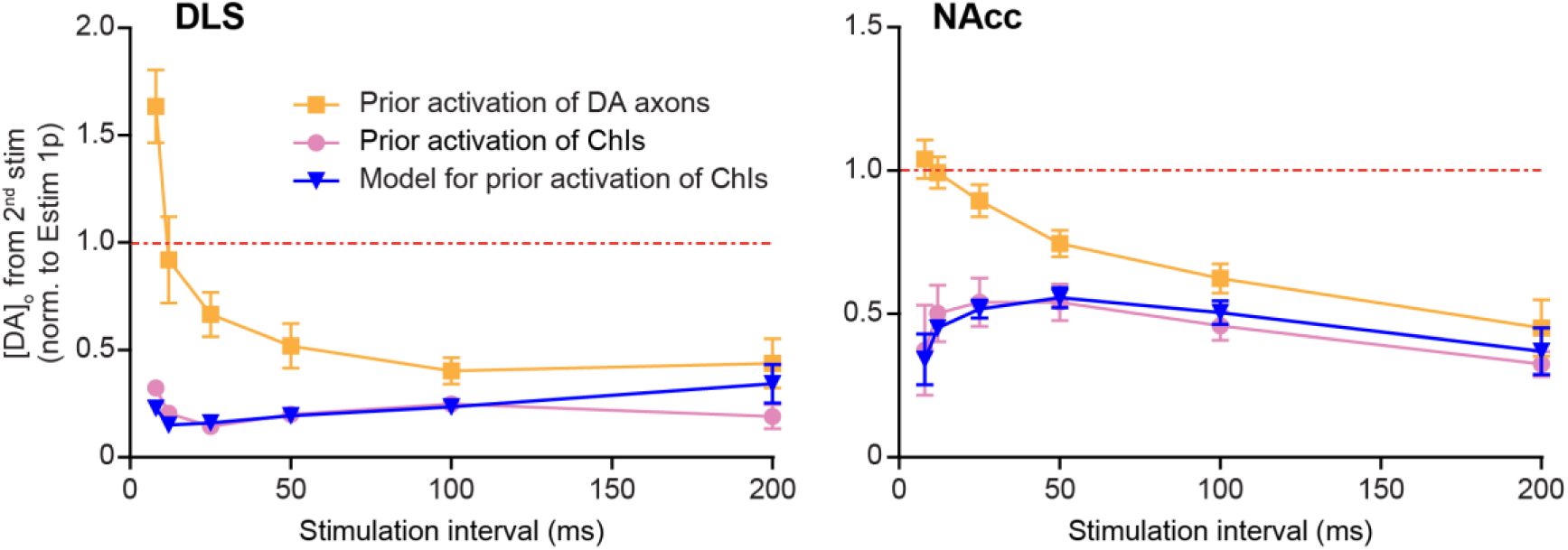
The computational model predicts empirical *ex vivo* data. The computational model built from empirical data in Fig. 3 D,H predicted the impact of prior activation of ChIs on subsequent DA release by an electrical stimulation as in Fig 1D,J. Because the electrical stimulation (the second stimulation in Fig 1d,h) evoked DA_DA_ and DA_ChI_ rather than only the DA_DA_ evoked by light stimulation (the second stimulation in Fig 3D,H), the predicted data (*blue*) was scaled as a whole curve to match the overall mean of the experimental data (*pink*) in DLS (*left*) and NAcc (*right*) to overcome the unknown ChI-induced inhibition of DA_ChI_ in this experimental paradigm.

**Extended Data Fig 7.**
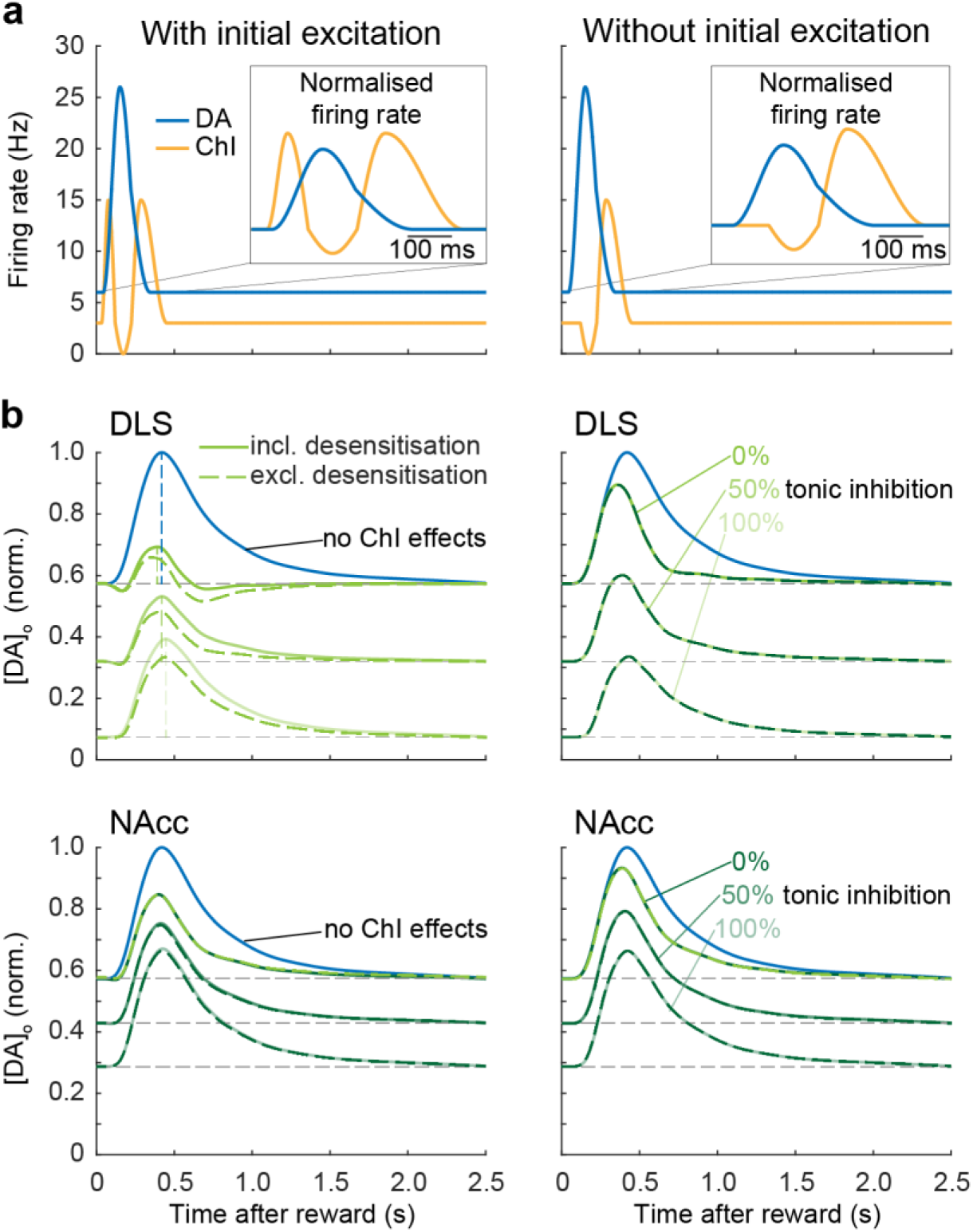
“Desensitisation” of nAChRs plays a limited role in modifying phasic DA release in a computational model during multiphasic ChI activity. **a**, Coincident phasic DA neuron firing (*blue*) and multiphasic ChI activity (*yellow*) with (*left*) or without (*right*) initial excitation in ChIs. Inset, firing rates normalised to baseline rates. **b**, Phasic DA release was modified by ChI activity (compared to no ChI effects, *blue*) both with (*solid green line*) and without (*dashed green line*) a “desensitisation” component in the model (obtained from Fig 4J,L) for DLS (*upper*) and NAcc (*lower*). Background levels of tonic inhibition of DA output by ChIs included in the model were 0, 50 and 100%. Inclusion of a dynamic desensitisation component in the model impacted on phasic DA release only when there was initial ChI excitation (*left)*, and only in DLS, acting to slightly increase phasic DA release by preventing nAChRs from inhibiting DA release during a rebound activity phase in ChIs. The effect is prominent in DLS (*upper*) than in NAcc (*lower*), consistent with the stronger ChI-induced inhibition of DA release in DLS than NAcc. As expected, excluding a dynamic desensitisation component from the model did not affect DA release when the initial excitation was absent because there was no initial increase in nAChR activation.

**Extended Data Fig 8.**
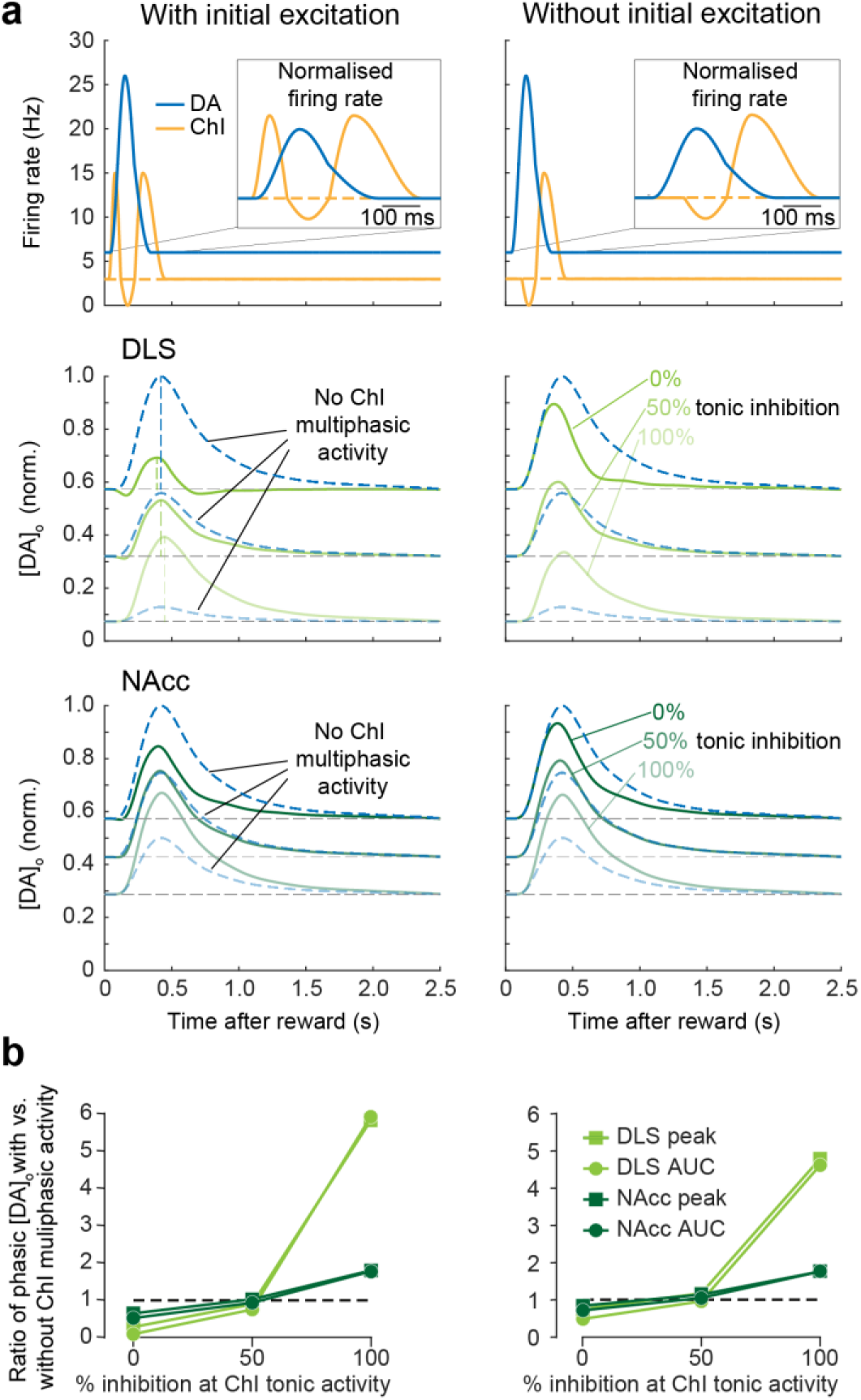
The tonic level of ChI-induced inhibition of DA release modifies the relative impact of ChI multiphasic activity on phasic DA release in a computational model. **a**, *Upper,* Model for coincident phasic DA neuron firing (*blue*) and multiphasic ChI activity (*yellow*) with (*left*) or without (*right*) initial excitation in ChIs. *Lower panels,* comparison of phasic DA signals at three different background levels of tonic inhibition of DA output (0, 50, 100%) before (*dashed blues*) versus after (*solid greens*) the development of multiphasic activity. At low-intermediate levels of tonic ChI inhibition of DA release (≤50%), multiphasic ChI activity reduced phasic DA release, whereas at very high levels of tonic ChI inhibition of DA release (50-100%), ChI multiphasic activity enhanced phasic DA release. **b**, Summary of impact of multiphasic ChI activity on phasic DA release on different background levels of tonic inhibition of DA release by tonic ChI activity. Change to phasic DA signal amplitude and area under curve (AUC) with (*left*) or without (*right*) a phase of initial ChI excitation.

## References

1 Schultz, W., Dayan, P. & Montague, P. R. A neural substrate of prediction and reward. Science 275, 1593–1599 (1997).

2 Rice, M. E. & Cragg, S. J. Nicotine amplifies reward-related dopamine signals in striatum. Nat Neurosci 7, 583–584 (2004). 10.1038/nn1244

3 Sulzer, D., Cragg, S. J. & Rice, M. E. Striatal dopamine neurotransmission: regulation of release and uptake. Basal Ganglia 6, 123–148 (2016). 10.1016/j.baga.2016.02.001

4 Threlfell, S. et al. Striatal dopamine release is triggered by synchronized activity in cholinergic interneurons. Neuron 75, 58–64 (2012). 10.1016/j.neuron.2012.04.038

5 Cachope, R. et al. Selective activation of cholinergic interneurons enhances accumbal phasic dopamine release: setting the tone for reward processing. Cell Rep 2, 33–41 (2012). 10.1016/j.celrep.2012.05.011

6 Mohebi, A. et al. Dissociable dopamine dynamics for learning and motivation. Nature 570, 65–70 (2019). 10.1038/s41586-019-1235-y

7 Mohebi, A., Collins, V. L. & Berke, J. D. Accumbens cholinergic interneurons dynamically promote dopamine release and enable motivation. Elife 12, e85011 (2023). 10.7554/eLife.85011

8 Liu, C. et al. An action potential initiation mechanism in distal axons for the control of dopamine release. Science 375, 1378–1385 (2022). 10.1126/science.abn0532

9 Wang, L. et al. Modulation of dopamine release in the striatum by physiologically relevant levels of nicotine. Nat Commun 5, 3925 (2014). 10.1038/ncomms4925

10 Wang, L. et al. Temporal components of cholinergic terminal to dopaminergic terminal transmission in dorsal striatum slices of mice. J Physiol 592, 3559–3576 (2014). 10.1113/jphysiol.2014.271825

11 Condon, M. D. et al. Plasticity in striatal dopamine release is governed by release-independent depression and the dopamine transporter. Nat Commun 10, 4263 (2019). 10.1038/s41467-019-12264-9

12 Ciesielska, A. et al. Anterograde axonal transport of AAV2-GDNF in rat basal ganglia. Mol Ther 19, 922–927 (2011). 10.1038/mt.2010.248

13 Brimblecombe, K. R. et al. Calbindin-D28K Limits Dopamine Release in Ventral but Not Dorsal Striatum by Regulating Ca2+ Availability and Dopamine Transporter Function. ACS chemical neuroscience 10, 3419–3426 (2019). 10.1021/acschemneuro.9b00325 PMID - 31361457

14 Condon, M. D. et al. Plasticity in striatal dopamine release is governed by release-independent depression and the dopamine transporter. Nature Communications 10, 4263 (2019). 10.1038/s41467-019-12264-9 PMID - 31537790

15 Katherine, R. B., Caitlin, J. G., Nicola, J. P. & Stephanie, J. C. Gating of dopamine transmission by calcium and axonal N-, Q-, T- and L-type voltage-gated calcium channels differs between striatal domains. The Journal of Physiology 593, 929–946 (2015). 10.1113/jphysiol.2014.285890 PMID - 25533038

16 Kramer, P. F., Twedell, E. L., Shin, J. H., Zhang, R. & Khaliq, Z. M. Axonal mechanisms mediating gamma-aminobutyric acid receptor type A (GABA-A) inhibition of striatal dopamine release. Elife 9, e55729 (2020). 10.7554/eLife.55729

17 Threlfell, S. et al. Striatal muscarinic receptors promote activity dependence of dopamine transmission via distinct receptor subtypes on cholinergic interneurons in ventral versus dorsal striatum. J Neurosci 30, 3398–3408 (2010). 10.1523/JNEUROSCI.5620-09.2010

18 Johnson, S. L., Marcotti, W. & Kros, C. J. Increase in efficiency and reduction in Ca2+ dependence of exocytosis during development of mouse inner hair cells. J Physiol 563, 177–191 (2005). 10.1113/jphysiol.2004.074740

19 Villette, V. et al. Ultrafast Two-Photon Imaging of a High-Gain Voltage Indicator in Awake Behaving Mice. Cell 179, 1590–1608 e1523 (2019). 10.1016/j.cell.2019.11.004

20 Kramer, P. F. et al. Synaptic-like axo-axonal transmission from striatal cholinergic interneurons onto dopaminergic fibers. Neuron 110, 2949–2960 e2944 (2022). 10.1016/j.neuron.2022.07.011

21 Aosaki, T. et al. Responses of tonically active neurons in the primate’s striatum undergo systematic changes during behavioral sensorimotor conditioning. J Neurosci 14, 3969–3984 (1994).

22 Morris, G., Arkadir, D., Nevet, A., Vaadia, E. & Bergman, H. Coincident but distinct messages of midbrain dopamine and striatal tonically active neurons. Neuron 43, 133–143 (2004). 10.1016/j.neuron.2004.06.012

23 Dani, J. A. & Bertrand, D. Nicotinic acetylcholine receptors and nicotinic cholinergic mechanisms of the central nervous system. Annu Rev Pharmacol Toxicol 47, 699–729 (2007). 10.1146/annurev.pharmtox.47.120505.105214

24 Changeux, J. P., Devillers-Thiery, A. & Chemouilli, P. Acetylcholine receptor: an allosteric protein. Science 225, 1335–1345 (1984). 10.1126/science.6382611

25 Giniatullin, R., Nistri, A. & Yakel, J. L. Desensitization of nicotinic ACh receptors: shaping cholinergic signaling. Trends Neurosci 28, 371–378 (2005). 10.1016/j.tins.2005.04.009

26 Shin, J. H., Adrover, M. F. & Alvarez, V. A. Distinctive Modulation of Dopamine Release in the Nucleus Accumbens Shell Mediated by Dopamine and Acetylcholine Receptors. J Neurosci 37, 11166–11180 (2017). 10.1523/JNEUROSCI.0596-17.2017

27 Exley, R. et al. Distinct contributions of nicotinic acetylcholine receptor subunit α4 and subunit α6 to the reinforcing effects of nicotine. Proceedings of the National Academy of Sciences 108, 7577–7582 (2011). 10.1073/pnas.1103000108 PMID - 21502501

28 Exley, R., McIntosh, J. M., Marks, M. J., Maskos, U. & Cragg, S. J. Striatal alpha5 nicotinic receptor subunit regulates dopamine transmission in dorsal striatum. J Neurosci 32, 2352–2356 (2012). 10.1523/JNEUROSCI.4985-11.2012

29 Collins, A. L., Aitken, T. J., Greenfield, V. Y., Ostlund, S. B. & Wassum, K. M. Nucleus Accumbens Acetylcholine Receptors Modulate Dopamine and Motivation. Neuropsychopharmacology 41, 2830–2838 (2016). 10.1038/npp.2016.81

30 Sun, F. et al. Next-generation GRAB sensors for monitoring dopaminergic activity in vivo. Nat Methods 17, 1156–1166 (2020). 10.1038/s41592-020-00981-9

31 Cunningham, C. L., Gremel, C. M. & Groblewski, P. A. Drug-induced conditioned place preference and aversion in mice. Nat Protoc 1, 1662–1670 (2006). 10.1038/nprot.2006.279

32 Collins, A. L. et al. Nucleus Accumbens Cholinergic Interneurons Oppose Cue-Motivated Behavior. Biol Psychiat 86, 388–396 (2019). 10.1016/j.biopsych.2019.02.014

33 Aosaki, T., Graybiel, A. M. & Kimura, M. Effect of the nigrostriatal dopamine system on acquired neural responses in the striatum of behaving monkeys. Science 265, 412–415 (1994).

34 Kramer, P. F. et al. Fast synaptic-like axo-axonal transmission from striatal cholinergic interneurons onto dopaminergic fibers. bioRxiv, 2022.2003.2025.485828 (2022). 10.1101/2022.03.25.485828

35 Krok, A. C. et al. Intrinsic dopamine and acetylcholine dynamics in the striatum of mice. Nature 621, 543–549 (2023). 10.1038/s41586-023-05995-9

36 Chantranupong, L. et al. Dopamine and glutamate regulate striatal acetylcholine in decision-making. Nature 621, 577–585 (2023). 10.1038/s41586-023-06492-9

37 Azcorra, M. et al. Unique functional responses differentially map onto genetic subtypes of dopamine neurons. Nat Neurosci 26, 1762–1774 (2023). 10.1038/s41593-023-01401-9

38 Zhang, H. & Sulzer, D. Frequency-dependent modulation of dopamine release by nicotine. Nat Neurosci 7, 581–582 (2004). 10.1038/nn1243

39 Kosillo, P., Zhang, Y. F., Threlfell, S. & Cragg, S. J. Cortical Control of Striatal Dopamine Transmission via Striatal Cholinergic Interneurons. Cereb Cortex 26, 4160–4169 (2016). 10.1093/cercor/bhw252

40 Cragg, S. J. Meaningful silences: how dopamine listens to the ACh pause. Trends Neurosci 29, 125–131 (2006). 10.1016/j.tins.2006.01.003

41 Schultz, W. Getting formal with dopamine and reward. Neuron 36, 241–263 (2002).

42 Howe, M. W. & Dombeck, D. A. Rapid signalling in distinct dopaminergic axons during locomotion and reward. Nature 535, 505–510 (2016). 10.1038/nature18942

43 Willuhn, I., Burgeno, L. M., Groblewski, P. A. & Phillips, P. E. Excessive cocaine use results from decreased phasic dopamine signaling in the striatum. Nat Neurosci 17, 704–709 (2014). 10.1038/nn.3694

44 Cragg, S. J., Hille, C. J. & Greenfield, S. A. Functional domains in dorsal striatum of the nonhuman primate are defined by the dynamic behavior of dopamine. J Neurosci 22, 5705–5712 (2002). 20026534

45 Willuhn, I., Burgeno, L. M., Everitt, B. J. & Phillips, P. E. Hierarchical recruitment of phasic dopamine signaling in the striatum during the progression of cocaine use. Proc Natl Acad Sci U S A 109, 20703–20708 (2012). 10.1073/pnas.1213460109

46 Hamid, A. A., Frank, M. J. & Moore, C. I. Dopamine waves as a mechanism for spatiotemporal credit assignment. bioRxiv, 729640 (2019). 10.1101/729640

47 Zhang, Y.-F., Fisher, S. D., Oswald, M., Wickens, J. R. & Reynolds, J. N. J. Coincidence of cholinergic pauses, dopaminergic activation and depolarization drives synaptic plasticity in the striatum. bioRxiv, 803536 (2019). 10.1101/803536

48 Zhang, Y. F., Reynolds, J. N. J. & Cragg, S. J. Pauses in Cholinergic Interneuron Activity Are Driven by Excitatory Input and Delayed Rectification, with Dopamine Modulation. Neuron 98, 918–925 e913 (2018). 10.1016/j.neuron.2018.04.027

